# Antigenic cancer persister cells survive direct T cell attack

**DOI:** 10.1101/2025.03.14.643359

**Authors:** Michael X. Wang, Brandon E. Mauch, August F. Williams, Tania Barazande-Pour, Filipe Araujo Hoffmann, Sophie H. Harris, Cooper P. Lathrop, Claire E. Turkal, Bryan S. Yung, Michelle H. Paw, David A. G. Gervasio, Tiffany Tran, Anna E. Stuhlfire, Theresa Guo, Gregory A. Daniels, Soo J. Park, J. Silvio Gutkind, Matthew J. Hangauer

## Abstract

Drug-tolerant persister cancer cells were first reported fifteen years ago as a quiescent, reversible cell state which tolerates unattenuated cytotoxic drug stress. It remains unknown whether a similar phenomenon contributes to immune evasion. Here we report a persister state which survives weeks of direct cytotoxic T lymphocyte (CTL) attack. In contrast to previously known immune evasion mechanisms that avoid immune attack, antigenic persister cells robustly activate CTLs which deliver Granzyme B, secrete IFNγ, and induce tryptophan starvation resulting in apoptosis initiation. Instead of dying, persister cells paradoxically leverage apoptotic caspase activity to avoid inflammatory death. Furthermore, persister cells acquire mutations and epigenetic changes which enable outgrowth of CTL-resistant cells. Persister cell features are enriched in inflamed tumors which regressed during immunotherapy *in vivo* and in surgically resected human melanoma tissue under immune stress *ex vivo*. These findings reveal a persister cell state which is a barrier to immune-mediated tumor clearance.

**Graphical abstract:** 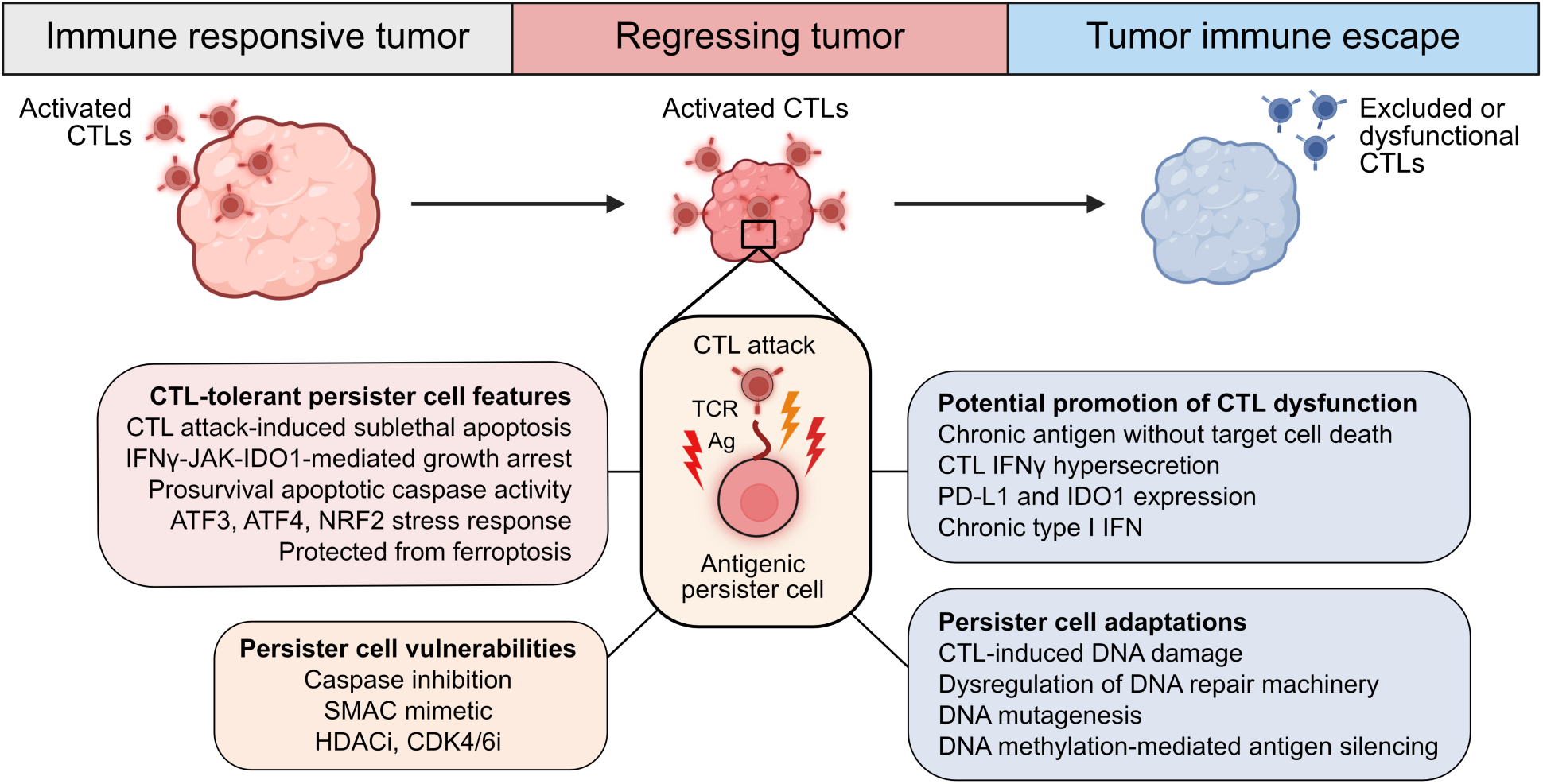

## Introduction

The ability of cells to enter into a persister state to endure cytotoxic drug stress was first observed in bacteria 80 years ago and in 2010 in cancer cells.^1–3^ Persister cells utilize reversible nongenetic mechanisms to survive drug treatment and contribute to residual disease and acquired drug resistance. Unlike drug resistant cells which engage stable mechanisms that avoid drug stress, persister cells leverage stress as demonstrated by recent reports that sublethal stress promotes persister cell formation,^4^ mutagenesis^5,6^ and regrowth during treatment (Williams A. F. et al., in revision^7^). Furthermore, sublethal drug stress in persister cells promotes emergent vulnerabilities, such as ferroptosis,^8^ which might be therapeutically targetable to eliminate residual disease and prevent acquired resistance.

Similar to the common failure of cancer drug treatments to induce durable complete responses, immunotherapy responses are also frequently incomplete.^9–11^ Indeed, partial responders are the most observed response group in advanced melanoma patients treated with dual immune checkpoint blockade (ICB) with nearly half of all responders relapsing within three years.^12^ Furthermore, instead of complete tumor clearance, many long-term survivors harbor residual disease.^12^ Indeed, incomplete immune-mediated tumor regressions have been observed for over a century including early studies with immunostimulatory toxins resulting in tumors that “*temporarily reduced in size*”, “*decreased one-half*”, or “*nearly disappeared*” before seeding fatal recurrences.^13^ Why are initial antitumor immune responses frequently incomplete?

The mechanisms that limit complete tumor rejection remain under active investigation and various models have been proposed including immune-induced dormancy, equilibrium, and intratumor antigenic and immunogenic heterogeneity.^14–20^ However, over the last fifteen years, observations of drug-tolerant persister cells have motivated the hypothesis that a population of persister cells may survive direct immune attack.^21^ Ideally, this question could be resolved by *in vivo* analysis within an authentic tumor immune microenvironment. However, to assess the central dynamic property which defines persister cells,^3^ reversible long-term tolerance, rather than avoidance, of direct CTL attack, would necessitate monitoring of individual antigenic cancer cells undergoing CTL attack over the course of multiple weeks *in vivo* which is not currently feasible. This technical hurdle has resulted in an impasse. To overcome this, we instead first turned to simplified long term CTL-tumor cell coculture models with frequent CTL replenishment to examine cancer cells undergoing direct CTL attack for multiple weeks without CTL exhaustion.

Using this approach, we discovered highly antigenic persister cells that survive weeks of activated CTL exposure. Contrary to immune evasion mechanisms that limit immune detection or activation,^22^ antigenic persister cells activate CTLs which deliver Granzyme B, induce cytotoxic IFNγ-dependent tryptophan starvation, and drive apoptotic signaling in persister cells. Rather than die, persister cells leverage sublethal activation of apoptotic caspases to avoid inflammatory death. Persister cells transcriptionally resemble and express stress response markers present in human and mouse tumors which have regressed on immunotherapy, in contrast to cancer cells within immunologically inactive tumors that are typically primary resistant to immunotherapy. Unexpectedly, we also found that CTL-tolerant persister cells are markedly distinct from drug-tolerant persister cells including strong resistance to ferroptosis. Finally, persister cells eventually give rise to heterogeneous CTL-resistant clones with both epigenetic and mutational adaptations. These findings implicate persister cells as a barrier to complete tumor clearance independent of immune dysfunction or loss of tumor recognition.

## Results

### Identification of antigenic CTL-tolerant cancer persister cells

We adapted CTL coculture models to allow for multiple weeks of coculture by replenishing CTLs every three days to limit CTL exhaustion (**Figure 1A**). Human T cell receptor (TCR) CD8^+^ T cells targeting the cancer testis antigen NY-ESO-1 (NY-ESO-1 CTLs) were generated by transducing primary human CD8^+^ T cells with TCR-expressing retrovirus as previously described.^23^ Three human BRAF^V600E^ melanoma cell lines that endogenously express NY-ESO-1 antigen (A375, SKMEL37, and MEL624) and, to minimize potentially confounding antigenic heterogeneity, three A375 subclones with different levels of stable NY-ESO-1 expression (A375-A, A375-B, and antigen-negative A375-C) were cocultured with NY-ESO-1 CTLs at varying CTL densities (**Figure S1A-G**). Sufficiently high CTL densities (CTL^high^) fully eliminated NY-ESO-1 expressing cancer cell populations within days of exposure (**Figure S1F-G** and **S1O**). Lower CTL densities (CTL^low^) also killed a large proportion of A375 and SKMEL37 cancer cells but spared residual cancer cells which survived for weeks as individual cells or small clusters, reminiscent of drug-tolerant cancer persister cells,^2^ before seeding sporadic outgrowth of escape colonies (ECs) (**Figures 1B** and **S2A**). CTL^low^ culture conditions therefore model residual cancer cell survival and outgrowth during long term CTL exposure.

**Figure 1.**
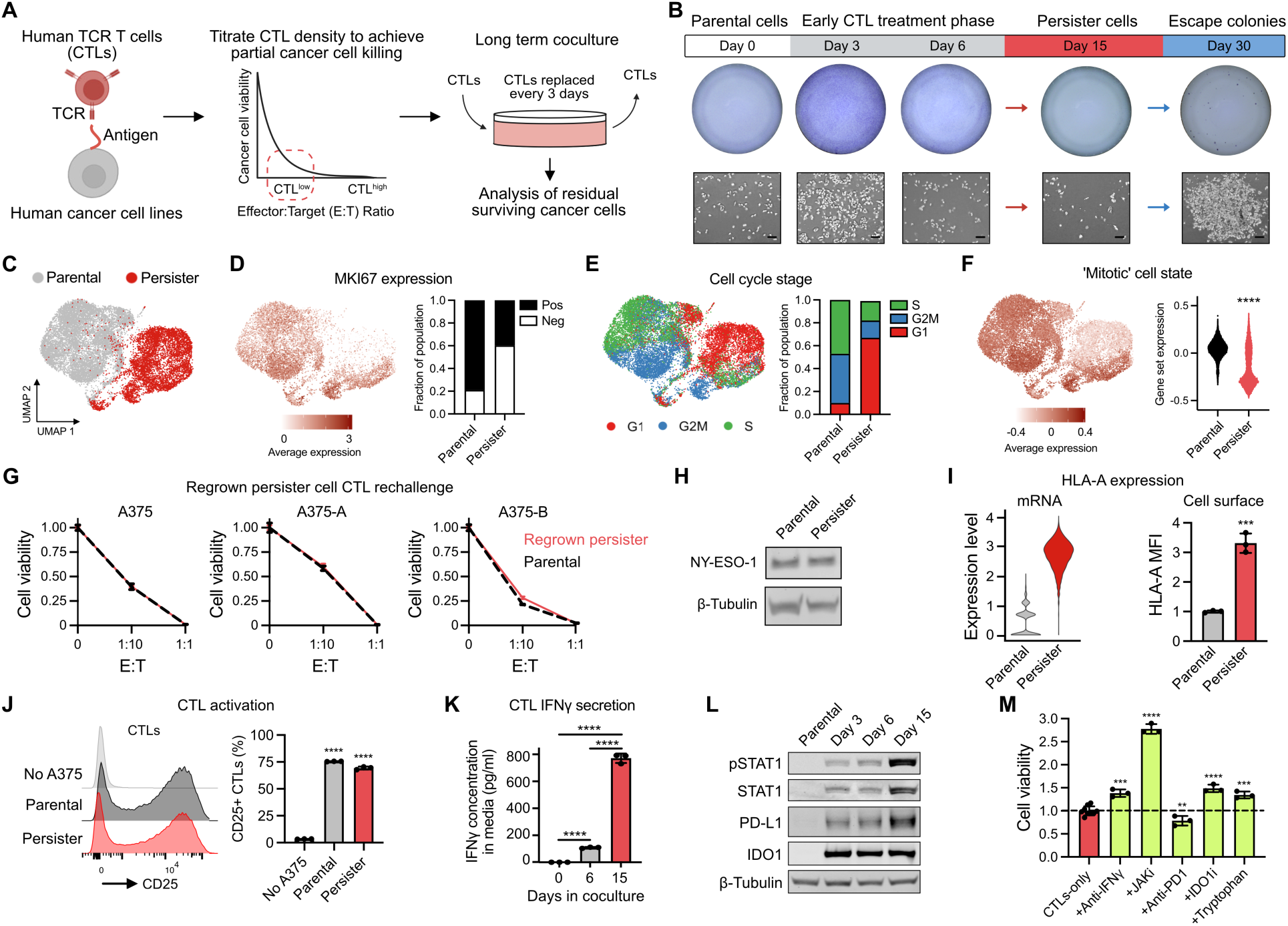
Antigenic persister cells survive weeks of activated CTL exposure. **(A)** Long term CTL coculture model. **(B)** A375 cells during NY-ESO-1 CTL^low^ coculture. Escape colonies are analyzed in Figure 5. Scale bars, 100 μm. (**C-F)** UMAP plot of parental and persister cell scRNA-seq **(C)**, proliferation marker MKI67 **(D),** and cell cycle stage analysis **(E)** in A375 scRNA-seq data. **(F)** Human melanoma tumor ‘mitotic’ signature^35^ in A375 cells (Mann-Whitney test). **(G)** A375 bulk and clonally derived A375-A and A375-B parental and persister cells allowed to regrow for one month without CTLs prior to rechallenge with NY-ESO-1 CTL^low^ coculture for 6 days. **(H)** Western blot of NY-ESO-1 expression in A375 cells. **(I)** Expression of HLA-A mRNA in scRNA-seq (left) and flow cytometry of cell surface HLA-A expression (right) in A375 cells (mean fluorescence intensity (MFI). **(J)** Flow cytometry analysis of CTL activation marker CD25 on CTLs cultured alone, with A375 parental cells for 3 days or persister cells for days 12-15 of coculture (t-tests versus CTLs cultured alone). **(K)** Measurement of secreted IFNγ concentrations in media collected during CTL^low^ coculture with A375 cells **(L)** Western blot of IFNγ-signaling nodes within A375 cells during CTL^low^ coculture. **(M)** A375 persister cell viability after 15 days of CTL^low^ coculture with treatments added on days 9-15 (CTLs-only n = 9, all other conditions n = 3); neutralizing anti-IFNγ antibody (1 μg/mL), JAK inhibitor (1 μM ruxolitinib), anti-PD-1 antibody (10 μg/mL pembrolizumab), IDO1 inhibitor (2 μM epacadostat), tryptophan (100 μg/mL). N = 3, mean ± SD are plotted, and two-tailed unpaired t-tests were performed unless stated otherwise. \**P* < 0.05; \*\**P* < 0.01; \*\*\**P* < 0.001; \*\*\*\**P* < 0.0001. See also **Figures S1** and **S2**.

To understand the cytotoxic stresses that residual cancer cells must survive, we next determined how outnumbered CTLs control and kill abundant cancer cell populations in coculture. CTL^low^ effector function against A375 and SKMEL37 cells was strongly reduced in the presence of neutralizing IFNγ antibodies, indoleamine 2,3-dioxygenase 1 (IDO1) inhibitor or excess tryptophan, but not neutralizing antibodies directed against TNF or FasL, or perforin inhibitor concanamycin A (**Figure S1H-I**). Recombinant IFNγ treatment also induced IDO1- and tryptophan-dependent growth arrest and cytotoxicity of A375 and SKMEL37 cells (**Figure S1J-L** and **S1O**), including NY-ESO-1 negative CTL-resistant subclone A375-C, indicating that antigen-loss variants can be controlled through bystander IFNγ signaling^24–26^ and tryptophan depletion. These observations are consistent with earlier reports that IDO1-mediated tryptophan depletion is a key mechanism of IFNγ-dependent tumor cell growth inhibition, an effect which has recently emerged as a potential contributor to the failure of IDO1 inhibitor clinical trials.^27–34^ Additionally, the pan-caspase inhibitor Q-VD-OPh (QVD), but not the ferroptosis inhibitor ferrostatin-1, partially protected A375 cells against CTL^low^ and IFNγ exposure, implicating caspase-dependent apoptosis but not ferroptosis in CTL^low^- and IFNγ-induced initial cancer cell death (**Figure S1M**).

Following two weeks of CTL^low^ coculture, single cell RNA sequencing (scRNA-seq) analysis revealed residual A375 cells are primarily growth arrested in the G1 cell cycle state, have reduced Ki-67 expression, and downregulate a mitotic cell state recently described in primary human melanoma^35^ (**Figure 1C-F**). Removal of CTLs after two weeks of coculture permitted the regrowth of the residual cancer cells which, following a one month holiday of regrowth without CTLs, became resensitized to CTLs (**Figures 1G** and **S2B**). Therefore, cancer cells can reversibly enter a quiescent, CTL-tolerant state fulfilling a key criteria of persister cells. We term the residual cancer cells that survive two weeks of CTL^low^ coculture as CTL-tolerant persister cells (CTL-persisters).

Persisters are notably antigenic, maintaining NY-ESO-1 antigen expression, increasing HLA-A mRNA and cell surface protein expression, and robustly activating CTLs resulting in high levels of CTL IFNγ secretion (**Figure 1H-K**). As a result, persisters exhibit chronically activated IFNγ-STAT1 signaling including STAT1 phosphorylation (pSTAT1) and induction of PD-L1 and IDO1 expression (**Figures 1L** and **S2C**). Importantly, the addition of neutralizing IFNγ antibody, JAK inhibitor, IDO1 inhibitor, or tryptophan supplementation promoted the outgrowth of A375 and SKMEL37 persisters (**Figures 1M, S1P,** and **S2D**). Therefore, chronic IFNγ-JAK-IDO1 signaling and tryptophan depletion enforce persister growth arrest. Furthermore, as expected because CTLs were replenished every three days to limit exhaustion, the addition of PD-1 blocking antibody had minimal effect on persister viability implicating survival mechanisms distinct from the PD-1/PD-L1 axis (**Figures 1M** and **S2D**). Together, these data indicate that persisters remain antigenic, activate CTLs and survive while undergoing chronic IFNγ signaling.

In contrast to A375 and SKMEL37 cells, MEL624 cells express low levels of NY-ESO-1, harbor defective IFNγ-STAT1 signaling, resist recombinant IFNγ, and are insensitive to NY-ESO-1 CTL^low^ coculture (**Figure S1A, S1F, S1H, S1J, and S1N-O**). Since MEL624 cells also express the melanoma differentiation antigen MART-1 (**Figure S3A**), we performed cocultures with MART-1 TCR CTLs. However, consistent with prior MART-1 TCR T cell melanoma studies,^36,37^ we observed dedifferentiation-driven rapid loss of MART-1 expression consistent with an early adaptive immune evasion mechanism rather than persistence (**Figure S3B-C**). Therefore, MEL624 largely avoid immune stress under CTL^low^ conditions. As a result, we instead focused on A375 and SKMEL37 NY-ESO-1 CTL^low^ coculture models to explore persister cells under immune stress.

### Persister cells survive direct CTL attack and undergo chronic sublethal apoptotic signaling

We found elevated amounts of intracellular granzyme B in live A375 persisters by flow cytometry indicating direct CTL attack without death (**Figure 2A**). Furthermore, live persisters exhibited chronic apoptotic signaling including cleaved caspase 3 and mitochondrial depolarization consistent with apoptotic mitochondrial outer membrane permeabilization (MOMP)^38,39^ (**Figure 2B-C**). We applied a fluorogenic apoptotic caspase 3/7 activity reporter to persisters and found that they exhibit elevated levels of caspase 3/7 activity above untreated cells but lower levels than cells treated with lethal concentrations of apoptosis inducer staurosporine (STS) (**Figures 2D** and **S4A**), indicating that persisters exhibit sublethal levels of apoptotic caspase activity. Consistent with sublethal apoptosis signaling, persisters with elevated caspase 3/7 activity could be sorted and replated without CTLs, remained viable and regrew (**Figure 2E**). ScRNA-seq analysis of persisters detected enrichment of genes previously shown to be upregulated in cells recovering from apoptotic stress,^40^ a process termed anastasis (**Figures 2F** and **S4B-C**), as well as upregulation of genes in the endosomal sorting complexes required for transport (ESCRT) pathway which was recently shown to protect cells from CTL attack by repairing perforin pores,^41^ further supporting that persisters survive rather than avoid CTL attack (**Figure S4D-E**).

**Figure 2.**
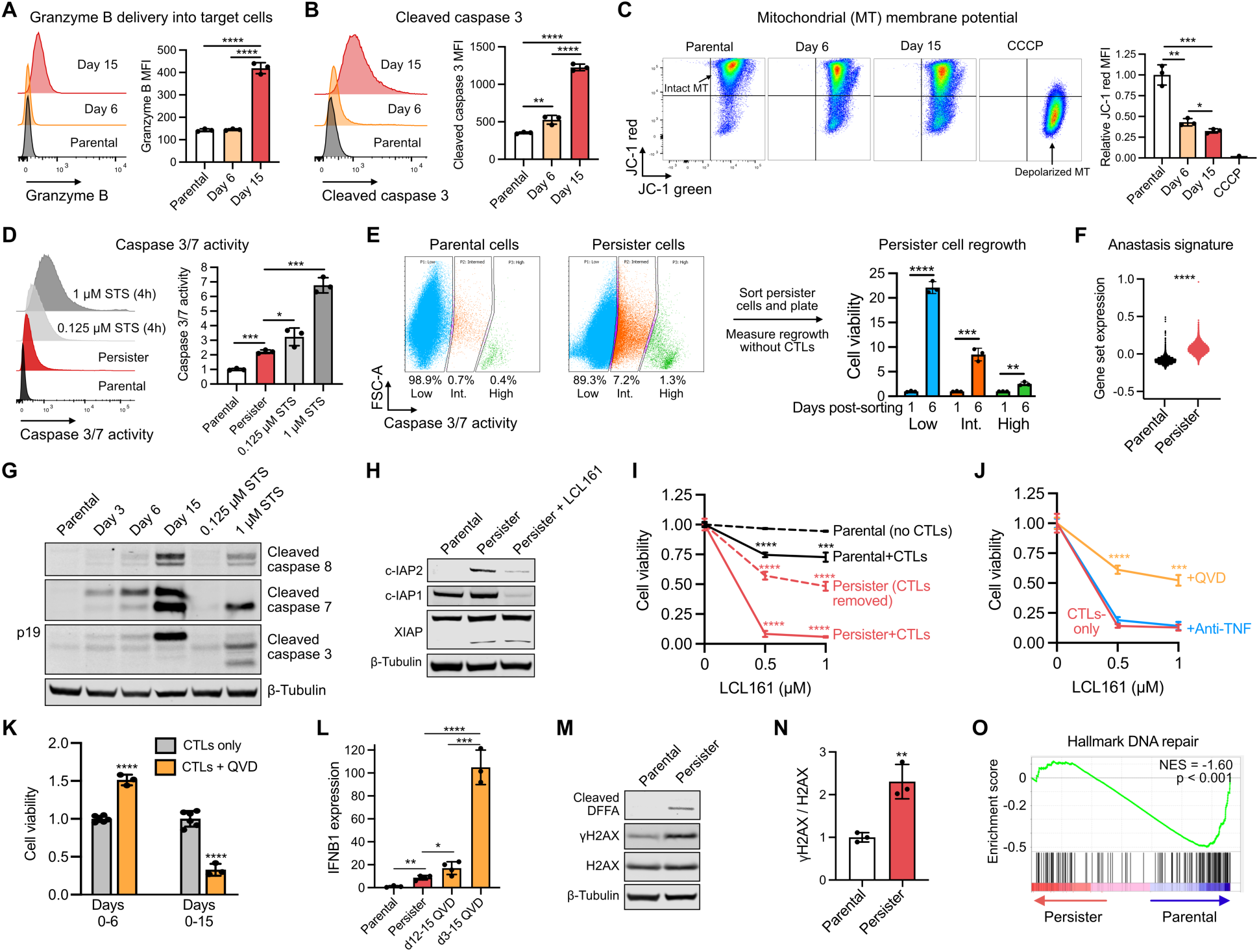
Persister cells survive CTL attack-induced apoptotic signaling. (A-O) Analysis of A375 cell CTL^low^ coculture. **(A-C)** Flow cytometry analyses of **(A)** granzyme B, **(B)** cleaved caspase 3 and **(C)** loss of mitochondrial (MT) polarity. **(D)** Flow cytometry of caspase 3/7 activity. **(E)** Cell viability of persister cells sorted for caspase 3/7 activity level and regrown without CTLs. **(F)** ScRNA-seq anastasis gene set^40^ signature score (Mann-Whitney test). **(G-H)** Western blots of **(G)** caspase cleavage and **(H)** IAPs including treatment with SMAC mimetic LCL161 (1 μM) for 15 hours. **(I)** Cell viability after LCL161 treatment in parental (3 days) or persister cells (days 12-15 of coculture) (t-tests versus parental). CTLs were absent or removed during LCL161 treatment where indicated. **(J)** A375 persister cells treated from days 12-15 with CTLs + LCL161 ± caspase inhibitor QVD (10 μM) or neutralizing TNF antibody (1 μg/mL) (t-tests versus CTLs-only). **(K)** Cell viability following treatment with CTLs ± 10 μM QVD (n = 6 for CTLs only, n = 3 for QVD, t-tests versus CTLs only). **(L)** IFNB1 mRNA expression (qRT-PCR) in parental cells, and in persister cells ± days 12-15 or 3-15 QVD co-treatment (n = 3-4). **(M)** Western blot of DFFA cleavage and DNA damage (γH2AX) and **(N)** quantification of γH2AX western blot signal. **(O)** Hallmark ‘DNA repair’ GSEA of differentially expressed genes in scRNA-seq data. N = 3, mean ± SD are plotted, and two-tailed unpaired t-tests were performed unless stated otherwise. \**P* < 0.05; \*\**P* < 0.01; \*\*\**P* < 0.001; \*\*\*\**P* < 0.0001. See also **Figures S2** and **S4**.

We also observed cleavage of caspase 8, caspase 7, and incomplete cleavage of caspase 3 resulting in persister accumulation of the caspase 3 p19 fragment, a marker of cells exposed to apoptotic stimuli during caspase inhibition (**Figures 2G** and **S2E**).^42–45^ Interestingly, levels of caspase cleavage, granzyme B delivery and IFNγ signaling (pSTAT1 and PDL1 upregulation), were higher in persisters at day 15 of coculture than in live cells earlier during coculture, indicating an accumulation of CTL-induced stress over time without death (**Figures 1L, 2A-B,** and **2G**). Based on these observations, we hypothesized that persisters survive CTL attack in part by inhibiting caspase activity.

### Caspase-active CTL-tolerant persister cells rely on IAP proteins to survive

We found that inhibitor of apoptosis (IAP) factors XIAP, c-IAP1 and c-IAP2 are increased within residual A375 persisters (**Figures 2H** and **S4F**) and that treatment of persisters with SMAC mimetic LCL161,^18,46^ which reduces persister c-IAP1 and c-IAP2 protein levels within hours (**Figure 2H**), eliminated ∼90% of persisters within three days (**Figure 2I**). Furthermore, when CTLs were removed from persisters, LCL161 treatment remained toxic (**Figure 2I**) demonstrating that the persister IAP dependency does not fully require simultaneous CTL attack. The tested LCL161 concentrations were nontoxic to parental cells and only modestly improved initial CTL-mediated killing of cancer cells (days 0-3 of coculture) (**Figure 2I**), indicating that SMAC mimetic targets an emergent vulnerability specific to persisters. SMAC mimetics have been previously reported to kill cells by lowering their TNF cytotoxicity threshold.^18,47^ However, neutralizing TNF antibody did not protect persisters from LCL161-induced death (**Figure 2J**). Instead, we found that the addition of the pan-caspase inhibitor QVD protected approximately half of CTL-exposed persisters from LCL161-induced death (**Figure 2J**), indicating LCL161-induced persister death is dependent on caspase activity. Therefore, although SMAC mimetics have failed to improve ICB responses in clinical trials^48^ potentially due to adversely interfering with immune function,^49^ CTL-tolerant caspase-active persisters rely upon IAPs to limit apoptosis which can be overcome by treatment with SMAC mimetic.

### Persister cells utilize sublethal apoptotic caspase activation to survive

Apoptotic caspases have additional functions beyond executing cell death.^38,50^ Persisters exhibit chronic caspase activation without death, and therefore we examined consequences of inhibiting caspase activity during coculture. While addition of QVD at the onset of CTL exposure (days 0-6) protected A375 cells from CTL-induced death, prolonged QVD treatment (days 0-15) markedly reduced the number of persisters (**Figure 2K**), demonstrating that chronic caspase activity paradoxically promotes the survival of persisters. Caspase-independent cell death (CICD) in caspase-inhibited cells has been previously reported to result from progressive mitochondrial dysfunction resulting from MOMP,^51,52^ leading to inflammatory death. Caspase activity has also been demonstrated to restrict type I IFN production.^53–56^ Consistent with this role for caspases in persisters, we found that while persisters increased IFNB1 expression above parental cells, QVD exposure in persisters dramatically further increased IFNB1 expression (**Figure 2L**), suggesting that CTL-induced caspase activity limits type I IFN production in persisters. Since inhibition of IFN signaling with JAK inhibitor treatment promotes persister outgrowth (**Figure 1M**), elevated IFN signaling upon caspase inhibition may contribute to decreased persister viability. We also observed cleavage of DNA Fragmentation Factor A (DFFA) within persisters (**Figure 2M**), a caspase 3 target that upon cleavage releases active apoptotic DNase DNA Fragmentation Factor B (DFFB) which induces DNA damage.^7^ Persister DNA damage (**Figures 2M-N** and **S2E**) together with dysregulation of DNA repair genes (**Figure 2O**), including genes previously implicated in stress-induced mutagenesis^5^ (**Figure S4G**), may promote mutagenesis and tumor evolution. Therefore, CTL-tolerant persisters leverage residual caspase activity to survive and adapt, presenting a complex role of caspase activity during anti-tumor immune responses.

### Immunotherapy-treated regressed tumors are enriched for persister cell features

Analysis of transcriptional cell states recently identified within human melanoma tumors treated with ICB revealed that persisters strongly resemble an antigen presentation state enriched within immune-infiltrated tumor niches and tumors responding to ICB (**Figure 3A-B**).^35^ Persisters are also enriched for interferon αβ response and neural crest-like melanoma signatures which were observed to reside in cells in close spatial proximity to antigen presentation cells in human tumors, consistent with persisters occupying antigenic inflamed tumor niches.^35^ Resembling persisters, this antigen presentation cell state includes upregulated interferon stimulated genes (ISGs) including STAT1, IRF1, HLA-A, and WARS, is distinguished from other melanoma cell states by increased expression of c-IAP2 and contains more genes implicated in anastasis than any other melanoma cell state (**Figure 3C-E**). These upregulated genes are consistent with an antigenic IFNγ-responsive cell state with anti-apoptotic features, mirroring our *in vitro* observations of CTL- tolerant persisters and suggesting that persisters may constitute a subset of antigen presentation melanoma cells in human tumors.

**Figure 3.**
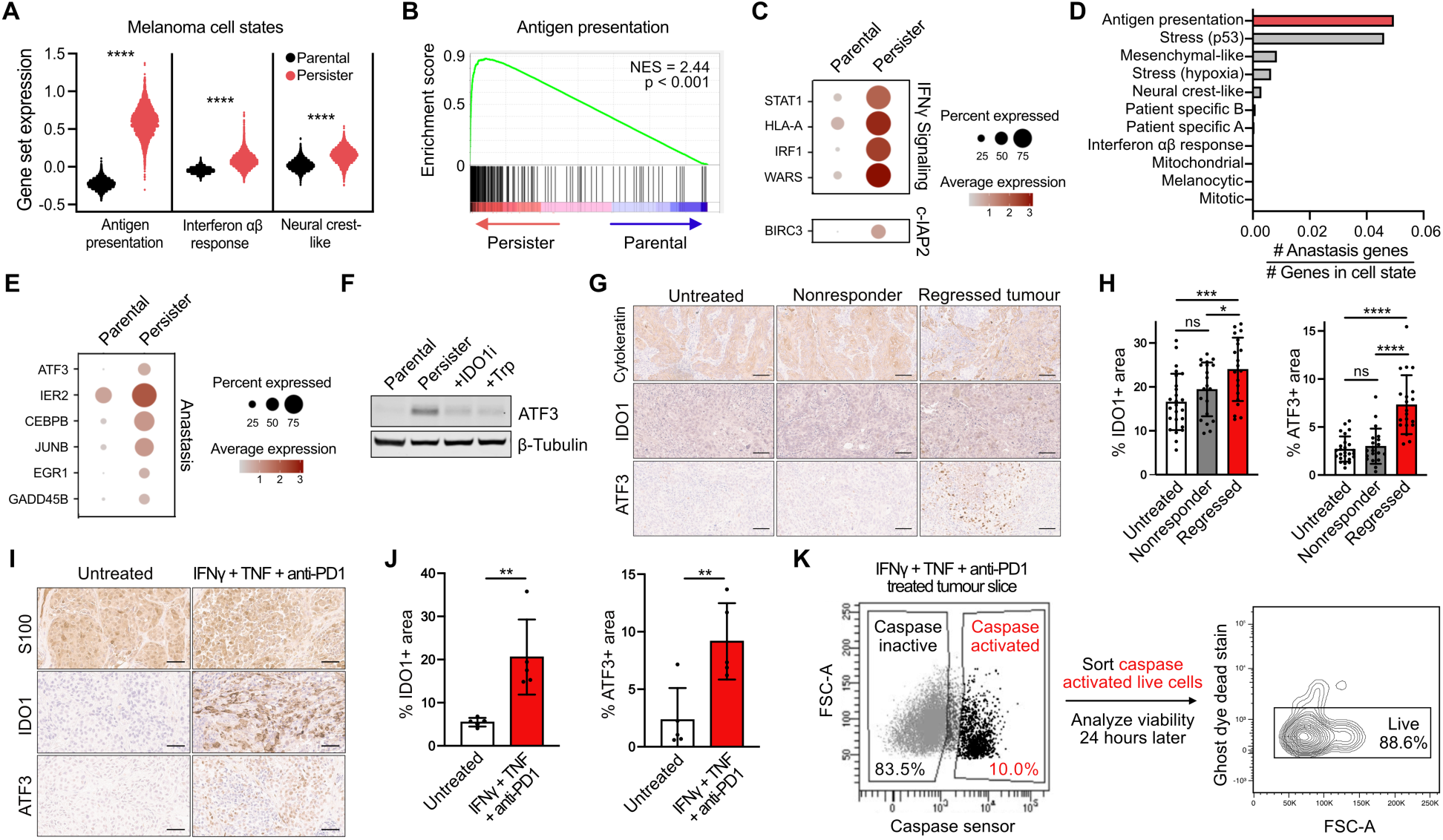
Persister cell features are enriched in immunotherapy-treated human and mouse tumors. A375 parental and CTL-tolerant persister cell scRNA-seq analysis of **(A)** signature scores of melanoma cell states observed in patients^35^ (Mann-Whitney test), **(B)** GSEA of the antigen presentation melanoma cell state and **(C)** expression of selected antigen presentation cell state genes. **(D)** Proportion of genes in each melanoma cell state which are anastasis-associated genes^40^ and **(E)** A375 cell scRNA-seq expression of selected anastasis-associated genes. **(F)** ATF3 expression in A375 persisters ± 2 μM IDO1 inhibitor epacadostat or 100 μg/mL tryptophan supplementation added during coculture days 12-15. **(G-H)** IHC analysis of 4MOSC1 head and neck squamous cell carcinoma syngeneic tumors from mice treated with 10 mg/kg anti-PD-1. **(G)** Representative IHC images and **(H)** quantification (n = 4-5 mice, 5 regions were analyzed per tumor). Scale bars, 50 µm. **(I-K)** Analysis of surgically resected primary human cutaneous melanoma tissue treated with 10 μg/mL anti-PD-1, 10 ng/mL IFNγ, and 10 ng/mL TNF for 6 days in culture. **(I)** Representative IHC images and **(J)** quantification (n = 1, five regions were analyzed per tumor slice). Scale bars, 50 µm. **(K)** Treated primary melanoma cells with elevated caspase 3/7 activity were sorted, replated, and tested for viability 24 hours later by flow cytometry (n = 1). Mean ± SD are plotted and two-tailed unpaired t-tests were performed unless stated otherwise. ns *P* > 0.05; \**P* < 0.05; \*\**P* < 0.01; \*\*\**P* < 0.001; \*\*\*\**P* < 0.0001. See also **Figure S5**.

A persister protein marker would facilitate detection of persisters but no broadly applicable marker has been identified. We and others have recently reported that the environmental stress response factor ATF3 is upregulated in drug-tolerant persisters,^4,7,57,58^ in cells recovering from apoptosis,^40^ and elevated ATF3 is included within the human melanoma antigen presentation cell state that otherwise resembles CTL-tolerant persisters (**Figure 3E**). ATF3 is also induced in CTL-tolerant persisters from stress resulting from IDO1-mediated tryptophan depletion (**Figures 3F** and **S2F**). Therefore, ATF3 is a broadly applicable persister marker. Consistent with persister enrichment in ICB treated tumors, ATF3 and IDO1 are induced in 4MOSC1^59^ orthotopic syngeneic mouse head and neck squamous cell carcinoma tumors which have regressed during anti-PD-1 treatment (**Figures 3G-H** and **S5A**). We also tested whether primary human tumors under immune stress exhibit persister features. Surgically resected human cutaneous melanoma tumor tissue was exposed to six days of immune stress by treating with anti-PD-1, IFNγ and TNF resulting in induction of IDO1 and ATF3 (**Figures 3I-J** and **S5B**). We then assessed a core functional feature of persisters, survival of apoptotic stress, in human tumor tissue. Consistent with sublethal apoptosis in immune stressed human tumors, treatment of melanoma tumor slices with anti-PD-1, IFNγ and TNF induced caspase-active yet viable tumor cells (**Figures 3K** and **S5B-D**). Therefore, immune-stressed human and mouse tumors contain multiple features indicative of CTL-tolerant persisters.

### Distinct cancer persister states survive CTL- and drug-induced stress

CTL-persisters share certain features with drug-persisters including reversibility, quiescence, sublethal apoptosis, and ATF3 induction. Beyond these features, it is unclear whether persisters which survive CTLs or drug treatment occupy a common survival state or are distinct. To test this, we generated CTL-persisters and drug-persisters from the same A375 parental cells, wherein drug-persisters were derived from BRAF and MEK inhibition, resulting in two persister populations with distinct morphologies (**Figures 4A** and **S6A**). We analyzed each population by scRNA-seq (**Figure 4B**) and, to examine transcriptional changes relevant to human tumors, we first assessed parental and persister expression of gene signatures within ICB-treated human melanomas^35^ (**Figures 4C** and **S7A-B**). Both persister populations downregulated the mitotic melanoma signature, consistent with their shared growth arrest phenotype. However, unlike drug-persisters which have reduced mitogenic signaling due to drug-induced suppression of phospho-ERK (pERK), CTL-persisters have elevated pERK (**Figure S6B**) indicating a distinct mechanism of growth arrest which we found is instead dependent on IFNγ-IDO1-mediated tryptophan depletion (**Figures 1M, S1P,** and **S6B**). Both persister populations contain a small subpopulation of mitotic cells^60^ and upregulate a neural crest-like signature consistent with dedifferentiated persister states. However, while CTL-persisters strongly resemble the antigen presentation melanoma signature observed within inflamed niches of ICB-responding tumors, drug-persisters instead upregulate the mesenchymal-like signature associated immune evasion and ICB nonresponsive tumors.

**Figure 4.**
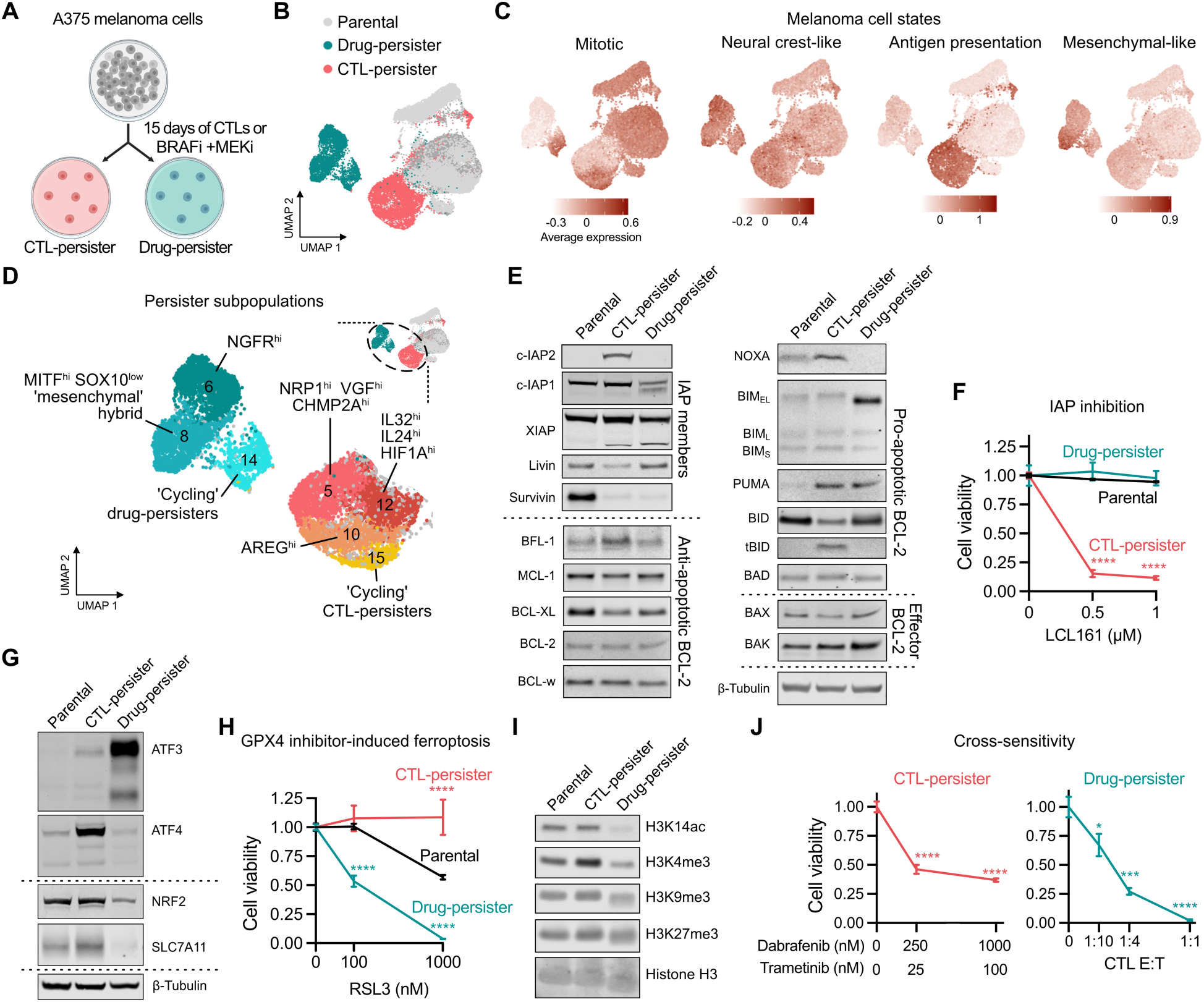
Cancer cells enter distinct persister states to survive CTL attack or drug stress. **(A)** Schematic of derivation of A375 persister cells that survive 15 days of NY-ESO-1 CTL^low^ coculture or 250 nM dabrafenib (BRAFi) and 25 nM trametinib (MEKi) treatment. **(B)** UMAP plot of A375 scRNA-seq. Dark and light grey are parental cells which were used for CTL and drug treatments, respectively. **(C)** UMAP plots highlighting expression of selected melanoma cell states.^35^ **(D)** UMAP plot subpopulations within drug-and CTL-persister populations. **(E)** Western blot analysis of IAP and BCL-2 family members. **(F)** Persister cell viability after 3 days of LCL161 treatment (n = 3, t-tests versus parental). **(G)** Western blot analysis of stress response and anti-ferroptosis factors. **(H)** Persister cell viability after 1 day of GPX4 inhibitor RSL3 treatment (n = 3-6, t-tests versus parental). **(I)** Western blot analysis of chromatin modifications in CTL- and drug-persisters. **(J)** CTL-persister viability after 3 day treatment with dabrafenib and trametinib (left) and drug-persister viability after 3 day coculture with CTLs (right). Initial CTL or drug exposure to derive persister cells was maintained during subsequent cotreatments. E:T is based on initially plated cell count prior to any treatment. (n = 3, t-tests versus untreated cells). Mean ± SD are plotted, and two-tailed unpaired t-tests were performed unless stated otherwise. \**P* < 0.05; \*\**P* < 0.01; \*\*\**P* < 0.001; \*\*\*\**P* < 0.0001. See also **Figures S6** and **S7**.

We next examined expression of recently described gene signatures that compose heterogeneity within human tumors, known as intratumor heterogeneity meta-programs (ITH MPs)^61^ (**Figure S7C**). CTL- and drug-persisters upregulate largely distinct ITH MPs, including nonoverlapping epithelial to mesenchymal transition (EMT) signatures, CTL-persister specific upregulation of stress and secretory signatures, and drug-persister specific upregulation of signatures identified within brain tumors (NPC/OPC, astrocytes). CTL- and drug-persisters also upregulate largely distinct hallmark gene sets, other than shared downregulation of cell cycle, DNA repair, and oxidative phosphorylation genes (**Figure S7D**). Finally, each persister state is composed of unique subpopulations including CTL-persister subpopulations defined by high Neuropilin 1 (NRP1) which may promote tumor vascularization^62^ and a cycling subpopulation which uniquely expresses EGFR ligand AREG^63,64^ (**Figures 4D** and **S7E**). Thus, CTL-persisters and drug-persisters occupy distinct transcriptional cell states.

Each persister state has similar levels of sublethal apoptotic caspase activity potentially reflecting the maximum sublethal threshold (**Figure S6C**). However, whether they arrive at this caspase activity level by adopting the same survival posture is unclear. To test this, we compared their levels of pro- and anti-apoptotic proteins and found that unlike the IAP-dependent survival state of CTL-persisters, increased IAP expression was not observed within drug-persisters which were correspondingly insensitive to SMAC mimetic (**Figure 4E-F**). There are also multiple differences in apoptotic machinery including that drug-persisters have absent NOXA and upregulated BIM while CTL-persisters have BFL1 upregulation, BID cleavage, and decreased BAX (**Figure 4E**), indicating distinct anti-apoptotic states. We also found that drug-persisters occupied an ATF3^very-high^, ATF4^low^, NRF2^low^ stress response state while CTL-persisters were characterized by a distinct ATF3^high^, ATF4^high^, NRF2^high^ state (**Figures 4G, S2F,** and **S7D**). While CTL-persister upregulation of ATF3 and ATF4^34^ is dependent on IDO1-mediated tryptophan depletion (**Figures 3F** and **S6D**), drug-persisters, which lack IFNγ signaling-mediated IDO1 expression, do not upregulate ATF4 and instead induce ATF3 through an alternative mechanism we recently reported.^7^

We previously found that drug-persisters are sensitized to death by ferroptosis.^8^ More recently, ferroptosis has been reported to play roles in primary responses to cancer immunotherapy.^65,66^ Surprisingly, we found that in contrast to drug-persisters, CTL-persisters are protected from rather than sensitized to ferroptosis (**Figures 4H** and **S2G**). Furthermore, we found that CTL-persisters upregulate NRF2 and its target SLC7A11, a component of the anti-ferroptosis cystine/glutamate antiporter system xc-, while drug-persisters downregulate SLC7A11, reflecting a ferroptosis-protective state in CTL-persisters (**Figures 4G** and **S2F**).^67^ This finding suggests that induction of ferroptosis may be ineffective in killing residual tumor cells which survive immunotherapy.

Beyond ferroptosis, we found that CTL- and drug-persisters possess other distinct vulnerabilities. Cotreatment with HDAC inhibitors reduced CTL- but not drug-persister viability, potentially reflecting their distinct chromatin states (**Figures 4I** and **S6E**). CTL-persisters also uniquely exhibited increased sensitivity to CDK4/6 inhibition^68^ (**Figure S6F**). Importantly, CTL-persisters were cross-sensitive to treatment with BRAF and MEK inhibitors, while drug-persisters were similarly sensitive to CTL exposure, indicating a lack of cross-resistance between persistent states (**Figures 4J** and **S6G**). Together, these data show that cancer cells adopt nonredundant persister states tailored to the stress encountered.

### Emergence of CTL-resistant escape clones from persister cells

When CTL-persisters were continually exposed to CTLs for one month, sporadic escape-colonies (ECs) emerged, including from A375-A and A375-B cells which were clonally derived to minimize preexisting heterogeneity (**Figures 5A** and **S8A**). We isolated 15 ECs from CTL- exposed A375 populations and 8 ECs from SKMEL37 cells, and each EC was expanded without CTLs. Isolated ECs exhibited varying growth rates after CTL removal and certain ECs exhibited distinct morphologies, reflecting the emergence of heterogeneous populations (**Figure S8B-C**). Contrary to the reversible CTL-tolerance of persisters (**Figures 1G** and **S2B**), most ECs exhibited stable resistance to CTLs upon reexposure after a CTL-free holiday and were less able to activate CTLs or elicit CTL IFNγ secretion (**Figures 5B-D** and **S8D**).

**Figure 5.**
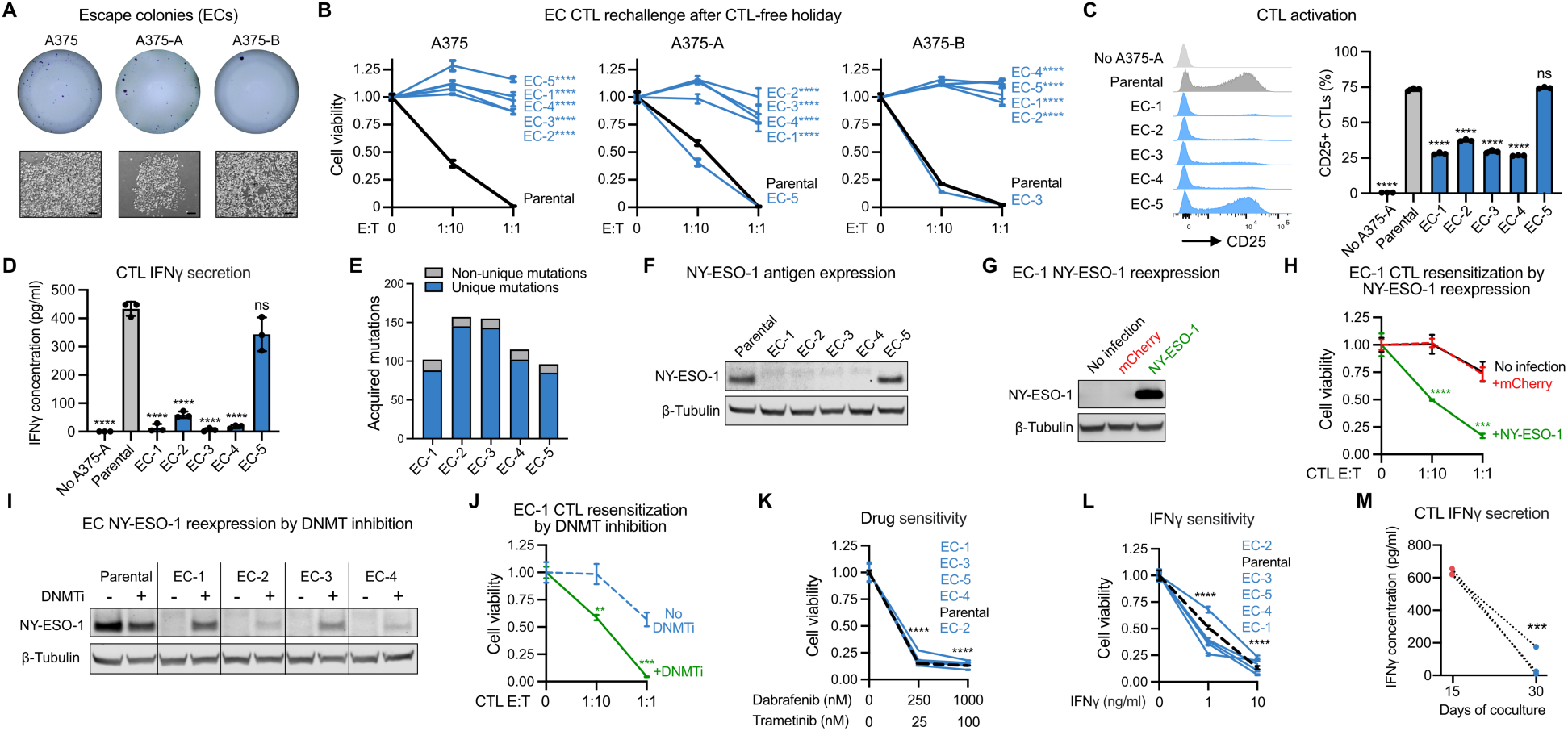
Heterogeneous escape colonies regrow from persister cells during CTL coculture. **(A)** ECs formed from bulk A375 and clonally derived A375-A and A375-B cell populations during 1 month of NY-ESO-1 CTL^low^ coculture. Scale bars, 100 μm. **(B)** ECs allowed to regrow without CTLs for one month before rechallenge with NY-ESO-1 CTL^low^ for 6 days (\*\*\*\**P* for both E:Ts versus parental). **(C-M)** A375-A cells and ECs previously regrown without CTLs were used unless otherwise stated. **(C)** Flow cytometry analysis of the activation marker CD25 on CTLs after 3 days of coculture (t-tests versus parental). **(D)** Soluble IFNγ after 3 days of CTL coculture (t-tests versus parental). **(E)** Acquired mutations identified within ECs. **(F)** Western blot of NY-ESO-1 expression in ECs and **(G)** EC-1 cells transduced to reexpress NY-ESO-1 or mCherry negative control. **(H)** Viability of NY-ESO-1 reexpressing EC-1 cells after 3 days of CTL coculture (t-tests versus no infection). **(I)** Western blot of antigen-loss ECs treated with the DNMT inhibitor (DNMTi) decitabine to reexpress NY-ESO-1 (1 μM, refreshed every day for 3 days). **(J)** EC-1 cell viability with 1 day pretreatment with 1 μM decitabine (DNMTi) followed by rechallenge with 3 days of NY-ESO-1 CTL coculture (t-tests versus no DNMTi treatment). **(K)** EC cell viability after 3 days of dabrafenib and trametinib treatment (\*\*\*\**P* for all cell lines and drug concentrations versus untreated cells) and **(L)** after 6 days of recombinant IFNγ exposure (n = 6 parental, n = 3 per EC, \*\*\*\**P* for all cell lines and IFNγ concentrations versus untreated cells). **(M)** Soluble IFNγ in media collected at days 15 and 30 of A375-A CTL^low^ coculture. N = 3, mean ± SD are plotted, and two-tailed unpaired t-tests were performed unless stated otherwise. ns *P* > 0.05; \**P* < 0.05; \*\**P* < 0.01; \*\*\**P* < 0.001; \*\*\*\**P* ≤ 0.0002. See also **Figure S8**.

Whole exome sequencing of A375-A derived ECs revealed approximately 100 unique mutations acquired by each EC during CTL exposure (**Figure 5E**). While mutagenesis may promote genetic immune-escape mechanisms over time, within this 1 month CTL exposure no putative resistance conferring mutations such as within antigen presentation or IFN pathway genes were identified (**Table S4**). Instead, we found that most A375 CTL-resistant ECs lost expression of the NY-ESO-1 antigen protein (**Figures 5F** and **S8E-F**) which was sufficient to avoid CTL activation because ectopic reexpression of NY-ESO-1 resensitized antigen loss ECs to CTLs (**Figure 5G-H**). Furthermore, treatment of antigen-loss ECs with the DNA methyltransferase inhibitor decitabine restored NY-ESO-1 expression and sensitivity to CTL exposure (**Figure 5I-J**), consistent with DNA methylation silencing of the NY-ESO-1 gene locus^69^ occurring within ECs. However, two of the thirteen CTL-resistant A375 ECs expressed NY-ESO-1 and three of the six CTL-resistant ECs isolated from SKMEL37 populations expressed comparable NY-ESO-1 levels to their parental population (**Figure S8E-G**), implicating additional immune evasion mechanisms beyond antigen loss. Also, all ECs remained sensitive to BRAF and MEK inhibitor treatment, indicating a lack of acquired cross-resistance to targeted oncogene-inhibition (**Figures 5K** and **S8H**).

Despite the ability of ECs to grow during CTL coculture, isolated ECs regrown without CTLs universally exhibited growth arrest or cell death when they were subsequently exposed to recombinant IFNγ (**Figures 5L** and **S8I**). IFNγ levels in coculture are highly elevated at day 15 when quiescent antigenic persisters but not ECs are present, but strongly decrease afterwards (**Figure 5M**). Given that inhibition of IFNγ signaling alone is sufficient to allow persisters to regrow (**Figure 1M**), these observations suggest that for ECs, whose properties are summarized in **Figure S8J**, outgrowth in coculture occurs as a direct result of decreased IFNγ levels allowing escape from IFNγ-enforced growth arrest. Therefore, IFNγ bystander signaling, which has been recently reported to influence tumor cells multiple cell layers beyond CTL-rich regions,^24–26^ is essential to maintain residual tumor cell quiescence including nonantigenic cells in coculture. We next asked whether IFNγ-enforced growth arrest can also be overcome by persister cells. To test this possibility, we exposed cancer cells to sustained immune effector cytokine levels.

### Persister cells can survive and escape cytotoxic immune effector cytokines

Chronic treatment of A375 cells with recombinant IFNγ produced residual IFNγ-tolerant persisters (cytokine-persisters) that, similar to CTL-persisters, exhibited intact IFNγ-signaling, IDO1- and tryptophan-dependent growth arrest, and ATF3 and ATF4 induction (**Figure 6A-E**). However, CTL- and cytokine-persisters exhibited differences including that while CTL-persisters have high levels of cleaved caspase 3, cytokine-persisters have low levels likely due to lack of granzyme B (**Figure 6C**). Notably the addition of the effector cytokine TNF, which alone is nontoxic to parental cells (**Figure 6F**), to IFNγ decreased the amount of cytokine-persisters which survived after 15 days (**Figure 6G**) yet suppressed persister IDO1 expression (**Figure 6H**) and promoted subsequent outgrowth of large colonies by day 30 (cytokine-ECs) (**Figure 6A-B** and **6I**), revealing intriguing dual roles for TNF in cytokine-persister behavior. Similar results were also observed with SKMEL37 cells (**Figure S9**). Interestingly, in contrast to predominantly irreversibly CTL resistant ECs isolated during coculture (**Figures 5B and S8D**), most isolated cytokine-ECs were resensitized to cytokine treatment following a cytokine-free holiday similar to regrown cytokine-persisters (**Figure 6J**). The rare cytokine-ECs that were stably resistant to cytokine reexposure (3/36 cytokine-ECs, **Figure 6J**) were also cross-resistant to CTL^low^ exposure and exclusively arose from heterogenous bulk A375 cell populations rather than clonally derived A375 cells (**Figure 6K**), consistent with selection of these rare irreversible cytokine-ECs from preexisting cytokine-resistant cells. These data indicate that, though rare cytokine-resistant cells may preexist, tumor cells can also engage reversible mechanisms to survive and regrow within IFNγ- and TNF-rich environments. Therefore, persister cells are a tumor cell subpopulation which can survive both direct CTL attack and bystander cytotoxic effector cytokine exposure.

**Figure 6.**
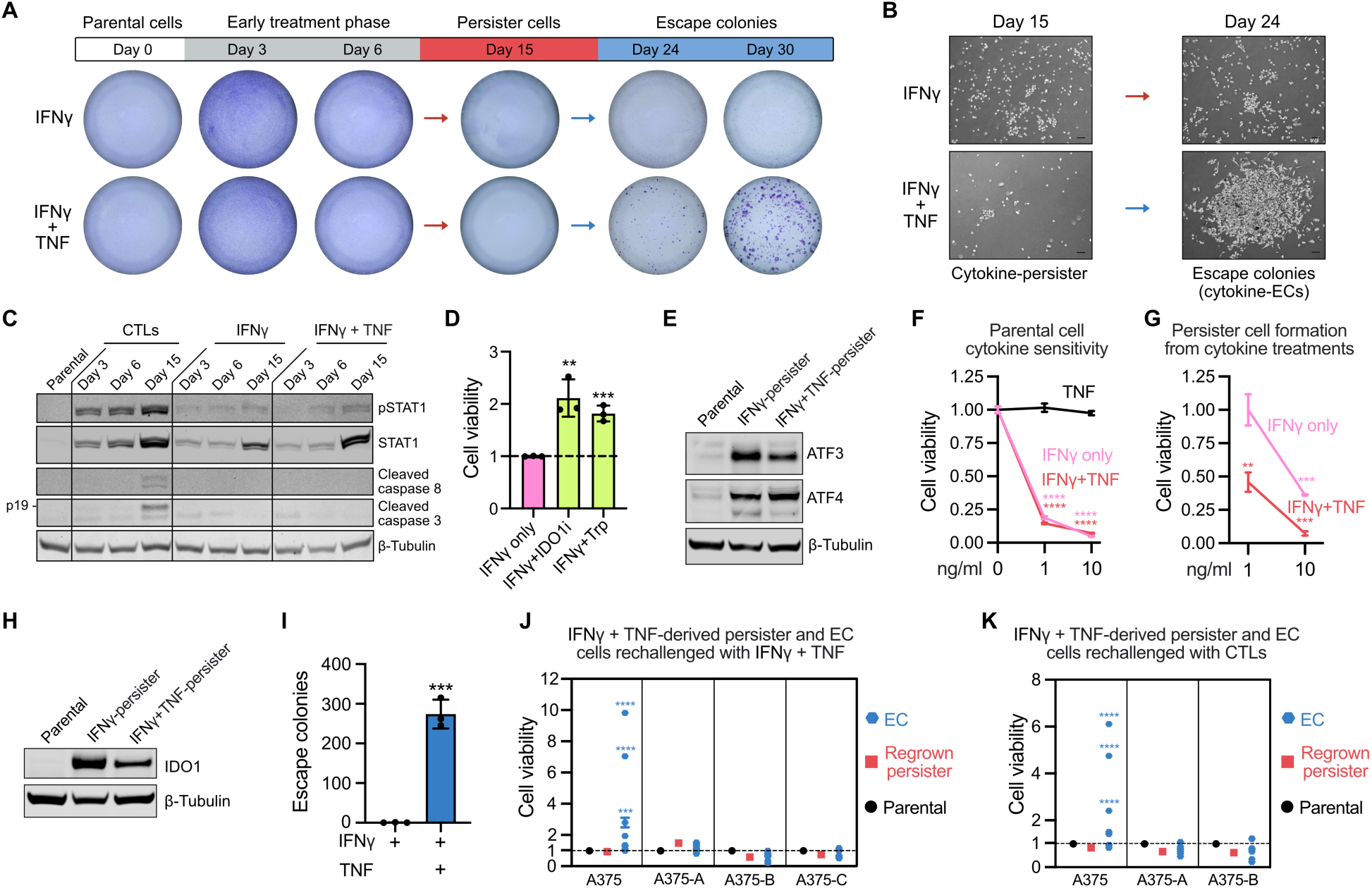
Cancer persistence against effector cytokines. **(A)** Representative crystal violet staining and **(B)** microscopy depicting A375-A cells during one month of recombinant IFNγ ± TNF exposure (1 ng/ml each). Scalebars, 100 μm. **(C)** Western blot of A375 persister cells which survived 15 days of CTL^low^ coculture or recombinant IFNγ ± TNF (1 ng/ml each) exposure. **(D)** A375 IFNγ-tolerant persister cells derived from 12 days of IFNγ exposure (1 ng/ml) were further treated from days 12-15 with IDO1 inhibitor (+IDO1i, 2 μM epacadostat) or tryptophan supplementation (+Trp, 100 μg/mL) with continued IFNγ exposure (t-tests versus IFNγ-only). **(E)** Western blot of ATF3 and ATF4 expression in A375 cytokine-persisters. **(F)** A375 parental cell viability after 6 days of cytokine treatment (t-tests versus untreated cells). **(G)** A375 cell viability after 15 days of recombinant IFNγ ± TNF exposure to form persister cells (t-tests versus 1 ng/ml IFNγ). **(H)** Western blot of IDO1 expression in A375 cytokine-persisters. **(I)** Quantification of A375-A EC formation after one month of IFNγ ± TNF exposure (1 ng/ml each). **(J)** Viability of A375 IFNγ + TNF-tolerant persister and EC cells regrown without cytokines and then rechallenged with 9 days of IFNγ + TNF exposure (1 ng/ml each) or **(K)** 6 days of CTL coculture (t-tests versus parental). The three cytokine-resistant ECs isolated from IFNγ ± TNF exposure of bulk A375 cells shown in **J** are the same ECs which are cross-resistant to CTL^low^ exposure shown in **K**. N = 3, mean ± SD are plotted, and two-tailed unpaired t-tests were performed unless stated otherwise. \**P* < 0.05; \*\**P* < 0.01; \*\*\**P* < 0.001; \*\*\*\**P* < 0.0001. See also **Figure S9**.

## Discussion

We found that human cancer persister cells remain antigenic, survive CTL attack and cytotoxic effector cytokine exposure, and seed outgrowth of CTL- and cytokine-resistant cells thereby implicating cancer persister cells as mediators of immune escape. Beyond merely surviving, persisters may promote immune dysfunction. For example, they serve as a source of chronic antigen (**Figure 1H**) which can exhaust or kill CTLs,^70^ promote CTL hypersecretion of IFNγ (**Figure 1K**) which may result from prolonged nonlethal CTL interactions,^71^ upregulate immune suppressive molecules such as PD-L1 and IDO1 (**Figure 1L**), and express chronic type I IFN (**Figure 2I**) which can drive immune dysfunction.^72–76^ Expression of PD-L1 additionally supports immune-independent cancer cell-intrinsic signaling that protects cancer cells from IFN cytotoxicity.^77,78^ Therefore, antigenic persisters appear equipped to both survive immune-induced cytotoxicity and drive immune-dysfunction, potentially playing a role in the transition from immunologically hot to cold tumors.

Furthermore, sublethal interactions between immune cells and persister cells may drive tumor genomic instability (**Figure 5E**) to generate nonimmunogenic clones that drive immune escape or, alternatively, generate neoantigens that sustain an immunologically active tumor, adding additional preclinical support that host immunity is an endogenous driver of tumor genomic instability.^79–81^ Indeed, when interpreting genomic analyses of clinical tumors that associate tumor mutation load with immune activity,^82^ one cannot exclude the alternative interpretation that immune activity induces mutagenesis of inflamed tumors. Further, whole exome sequencing of matched baseline and relapsed melanoma tumors treated with ICB suggest tumors may acquire genetic changes during immunotherapy.^83^ It is also plausible that persisters evolve under immune-induced stress without acquiring mutations. For example, CTL attack-induced DNA damage may induce DNA methylation events^84^ in persister cells that influence expression of critical genes involved in anti-tumor immunity, including tumor rejection antigens similar to NY-ESO-1 silencing that we observed (**Figures 5F** and **S8J**). Further, persisters that survive within IFNγ-rich tryptophan-depleted environments increase expression of the tryptophanyl-tRNA synthetase WARS (**Figure 3C**), raising the possibility that persisters utilize tryptophan-to-phenylalanine codon reassignments to influence protein activity and immunoreactivity.^85,86^ Therefore, the persister cell evolution model proposed in drug-treated bacteria and cancer cells^87^ is likely operational within stressed immunologically active tumors whereby sublethal interactions between immune cells and cancer cells drive malignant behavior, tumor instability, and neoantigen sculpting.

We also found that treatment with JAK or IDO1 inhibitors results in persister regrowth (**Figures 1M** and **S2D**), warranting caution when utilizing these inhibitors in cancer patients.^33,88,89^ In retrospect, the tumor cell-protective effect of IDO1 inhibition must be considered when reconciling the failure of IDO1 inhibitors to enhance ICB response in patients.^27–34^ As the immune oncology field explores the use of JAK inhibitors in the treatment of cancer, untangling the contextual pro- and anti-tumor effects of the IFN-JAK-STAT pathway(s) remains a pending challenge that requires re-evaluation of the roles of type I and II IFNs in antitumor immunity.^90^ Indeed a recent meta-analysis of CRISPR screens has highlighted a disparity in the role of IFNγ on antitumor immunity between coculture and *in vivo* tumor rejection models.^91^ In a similar vein as double-edged IFN-signaling, observations within our CTL- and cytokine-tolerant persister models implicate opposing pro- and anti-tumor roles of both apoptotic caspase activity (**Figure 2K**) and TNF signaling (**Figure 6**) on persister cell behavior that change over time, highlighting the importance of understanding divergent acute and chronic effects of these signaling nodes within antitumor immunity.

The identification of persister cell vulnerabilities presents rational targets for combination therapies. While we found that, unlike drug-tolerant persisters, CTL-tolerant persisters are resistant to ferroptosis, they are selectively sensitive to other treatments including dependence on apoptotic caspase activity for survival and therefore may be therapeutically targetable. As accumulating evidence implicate critical roles of caspases beyond the induction of apoptosis,^38,50,92–94^ inhibiting the pro-survival effects of caspase activity presents an intriguing avenue to improve antitumor immune responses. Furthermore, our study supports that CTL-tolerant persisters may utilize IAP activity (e.g., XIAP, c-IAP1, c-IAP2) to limit lethal caspase activity while under apoptotic stress,^95^ creating a persister vulnerability to SMAC mimetics in tumors which are under active CTL attack. In addition, persister cell engagement of ATF3, ATF4, and NRF2 stress responses, as well as occupying a surprising ferroptosis resistant cell state, may further contribute to their survival within immunologically active tumors and create vulnerabilities.

Beyond our findings, long-term coculture, recombinant effector cytokine, and tryptophan deprivation persister cell models will enable other investigators to explore immune-tolerant persister cells. Indeed, the curation of a scRNA-seq catalogue of persister cells across cancer types and therapies may be useful to incorporate into growing catalogues of human tumor and cell atlases.^96^ Together, these findings add another dimension to growing interest in how tumor cells survive immune attack^97^ by demonstrating that a subpopulation of CTL-sensitive antigenic cancer cells reversibly enter a quiescent persister state, undergo immune attack-induced apoptotic signaling yet survive.

### Limitations of the study

While coculture models have fueled the investigation of cellular immunity and cell-mediated cytotoxicity since 1960,^98^ it is reasonable to question the relevancy of these models to the *in vivo* setting. However, the core features of our models including CTL^low^ environments,^99^ sublethal immune-target interactions,^100,101^ and IFNγ bystander signaling^24–26^ have been observed *in vivo* within human or mouse tumors. Also, because persister cells are functionally defined rather than defined by markers, convincing demonstration that persister cells exist *in vivo* would conceivably require longitudinal intravital monitoring of individual tumor cell-CTL interactions for weeks which is not currently technically possible. However, the strong resemblance of the CTL-tolerant persister transcriptome to an antigenic cell state recently observed in responding human melanoma tumors treated with immunotherapy (**Figure 3A-E**), enrichment for the persister marker ATF3 in immune-mediated regressed human and mouse tumors (**Figure 3G-J**), and the observation that immune stress induces sublethal caspase activity within intact human melanoma tissue (**Figure 3K**) support the *in vivo* relevance of our findings. Last, we cannot exclude the possibility that the antigen loss escape variants isolated from persister cells were rare subclones that emerged prior to CTL exposure, despite generating single-cell derived A375-A and A375-B lines to combat this possibility.

## Resource availability

### Lead contact

Requests for further information and resources should be directed to and will be fulfilled by the lead contact, Matthew J. Hangauer.

### Materials availability

All unique plasmids and cell lines generated in this study are available from the lead contact without restriction.

## Method details

### Cancer cell lines

The human BRAF^V600E^ melanoma cell lines A375 (ATCC), SKMEL37 (Sigma), and MEL624 (provided by Nicholas Restifo and Steven Rosenberg, NCI) were cultured in RPMI 1640 medium (Gibco #11875119) supplemented with 10% FBS (Gibco #A5256801) and 1% antibiotic-antimycotic (Gibco #15240062) in a 5% CO_2_ atmosphere at 37°C, unless otherwise noted. Cell line identities were confirmed by short tandem repeat profiling by the UC Berkeley DNA Sequencing Facility and were found to be negative for mycoplasma using the MycoAlert kit (Lonza). To derive clonal A375 cell lines A375-A, A375-B, and A375-C, single cells were plated into 96 well cell culture plates, expanded into a 10 cm cell culture dish over 4 weeks, and frozen. The 4MOSC1 syngeneic mouse head and neck squamous cell carcinoma (HNSCC) cell line was developed and described for use in immunotherapy studies in prior reports.^59,102,103^ 4MOSC1 cells were cultured in Defined Keratinocyte-SFM medium (Gibco #10744019) supplemented with EGF Recombinant Mouse Protein (5 ng/ml, ThermoFisher #PMG8041), Cholera Toxin (50 pM, Sigma #C8052) and 1% antibiotic-antimycotic solution in a 5% CO_2_ atmosphere at 37°C.

### TCR CD8^+^ T cell production

We modified a previously described protocol^23^ to generate cytotoxic CD8^+^ T lymphocytes (CTLs) that recognize the HLA-A*02–restricted NY-ESO-1 antigen (NY-ESO-1:157–165 epitope) or MART-1 antigen (MART-1:27–35 epitope). Retroviral supernatants were collected from PG13 cells that express a retroviral vector for the NY-ESO-1 TCR or MART-1 TCR (provided by Nicholas Restifo and Steven Rosenberg, NCI) and frozen at −80°C. CD8^+^ T cells were isolated from frozen human PBMCs (STEMCELL #70025 or 70025.2) using the EasySep Human CD8^+^ T Cell Isolation Kit (STEMCELL #17953) per product protocol without the DNase I step. HLA-matching of PBMC donors and cancer cell lines to avoid alloreactivity was not performed because uninfected CTL alloreactivity was found to be low at E:T ratios less than 1:1 (**Figure S1E**), E:T ratios used for all persister cell experiments were 1:4 or lower, and most experiments were performed multiple times with CTLs derived from at least two different PBMC donors with consistent results in all cases. Isolated CD8^+^ T cells were incubated in ImmunoCult-XF T Cell Expansion Medium (XF-TCM, STEMCELL #10981) supplemented with 7.5 ng/mL human recombinant IL-2 (STEMCELL #78036.3) and 25 μL/mL ImmunoCult Human CD3/CD28/CD2 T Cell Activator (STEMCELL #10970) in a 5% CO_2_ atmosphere at 37°C for 48-72 hours prior to retroviral transduction. We hereafter refer to CD8^+^ T cells as CTLs.

To engineer CTLs to express TCRs, RetroNectin-assisted retroviral-mediated gene transduction was performed. Wells of 6 well non-treated cell culture plates (Genesee #25-100) were coated with 2 mL of 10 μg/mL RetroNectin (Takara #T100A) diluted in PBS at 4°C overnight. The next day, RetroNectin solution was removed from the cell culture plates, 4 mL of thawed TCR retroviral supernatant was added to each RetroNectin-coated well, and plates were centrifuged at 1,000g for 2 hours at 32°C to bind virus particles to RetroNectin reagent. After centrifugation, 3 mL of retroviral supernatant was removed from each well and 1 million CTLs were added to each well by adding 4 mL of 250,000 CTLs/mL in XF-TCM + 7.5 ng/mL IL-2. CTLs were incubated in RetroNectin-coated plates overnight in a 5% CO_2_ atmosphere at 37°C. The next day, a second round of transduction was performed in the same manner and CTLs were incubated for 24-48 hours prior to fluorescence-activated cell sorting (FACS) for TCR expression.

To isolate TCR expressing CTLs, CTLs were collected from RetroNectin-coated plates, washed with PBS, washed with PBS + 5% FBS, and incubated in 50 μL PBS + 5% FBS with 10 μL NY-ESO-1 Dextramer-PE (Immudex #WB2696-PE) or MART-1 Dextramer-PE (Immudex #WB2162-PE) per 1 million CTLs for 10 minutes at room temperature. CTLs were then washed with PBS + 1% FBS, incubated with a CD8-APC antibody (STEMCELL #60022AZ.1, diluted in PBS + 1% FBS) for 20 minutes at 4°C, washed 2x with PBS + 1% FBS, and resuspended at 25 million cells/mL in cold PBS for FACS using a BD FACSAria II High Speed Sorter. CD8 and TCR double positive cells were sorted into XF-TCM. After sorting, isolated CTLs were resuspended at 250,000 cells/mL in XF-TCM + 7.5 ng/mL IL-2 and incubated for 4-5 days in a 5% CO_2_ atmosphere at 37°C prior to expansion, maintaining CTL density between 100,000-1M cells/mL. To expand TCR CTLs, CTLs were resuspended at 1 million cells/mL in XF-TCM + 7.5 ng/mL IL-2 + 1.6 μL/mL ImmunoCult Human CD3/CD28/CD2 T Cell Activator and incubated for 2-3 days. CTL expansion was monitored an additional 5 days, maintaining CTL density between 100,000-600,000 cells/mL. To freeze and store TCR CTLs, CTLs were pelleted and resuspended in cold CrytoStor CS10 (STEMCELL #07959 or Sigma #C2874) at 10 million cells/mL and 1-2 million cells were aliquoted into single-use thaws, frozen overnight at −80°C in a Mr. Frosty Freezing Container (Nalgene #5100-0001), and transferred to liquid nitrogen for long-term storage. TCR CTLs were thawed 1-2 days prior to coculture. CTL activity varied by PBMC donor batch and therefore CTL densities were titrated specifically for each batch to induce effective but incomplete elimination of melanoma cell populations (CTL^low^ E:Ts ranged from 1:13 to 1:4).

### Persister cell models

Cancer cells were exposed to CTLs, recombinant cytokines, or drug in 10 cm cell culture dishes or 12 well cell culture plates in RPMI 1640 medium supplemented with 10% FBS and antibiotic-antimycotic in a 5% CO_2_ atmosphere at 37°C. Cancer cells were seeded at 500,000 cells per 10 cm dish or 20,000 cells per 12 well and were exposed to CTLs, cytokines, or drug the next day. 10 mL and 625 μL of media were used for 10 cm and 12 well cultures, respectively. Media with fresh CTLs, cytokines, or drug were replenished every 3 days for up to 30 days. To limit signals from detached cells, cell debris, and CTLs, plates were washed at least 3x with PBS or media prior to downstream analysis.

For CTL coculture, 1-2 days prior to coculture with cancer cells, CTLs were thawed and suspended at 500,000 CTLs/mL in ImmunoCult-XF T Cell Expansion Medium (STEMCELL #10981) supplemented with 7.5 ng/mL human recombinant IL-2 (STEMCELL #78036.3). CTLs were added to RPMI + 7.5 ng/mL IL-2 at the indicated effector to target ratios (E:T), and added to adhered cancer cells. Throughout all coculture experiments, CTLs were replenished every 3 days in fresh media with 7.5 ng/mL IL-2 at a constant CTL density matching the initial E:T. Residual cancer cells that survived 12-15 days of CTL exposure were classified as CTL-tolerant persister cells. Emerging cancer cell colonies were sporadically observed during CTL coculture and classified as escape colonies (ECs). In all cases, cancer cells were analyzed after removal of the nonadherent CTLs by rinsing the plates with media or PBS.

For effector cytokine exposure, cancer cells were exposed to 1-10 ng/mL of recombinant human IFNγ (Peprotech #300-02) ± TNF (Peprotech #300-01A). Cytokines and media were replenished every 3 days in all experiments. Residual cancer cells that survived 12-15 days of cytokine exposure were classified as cytokine-tolerant persister cells. Emerging cancer cell colonies observed during IFNγ + TNF exposure were classified as cytokine-escape colonies (cytokine-ECs).

For drug exposure, A375 cells were treated with 250 nM dabrafenib (Selleckchem #S2807) and 25 nM trametinib (Selleckchem #S2673) for 12-15 days. Drug and media were replenished every 3 days. Residual cancer cells that survived 12-15 days of drug exposure were classified as drug-tolerant persister cells. Unless otherwise stated, all persister cell experiments were performed on prederived persister cells while maintaining exposure to the CTLs, cytokines or drugs used to derive the persister cells.

### Regrown persister cell and escape colony cell lines

To generate regrown persister cells, CTLs or IFNγ and TNF were removed from residual persister cells that survived 15 days of coculture or cytokine exposure, plates were washed 3x with media, and persister cells were expanded for ∼1 month without CTL or cytokine exposure. The resulting populations were considered regrown persister cells.

To isolate escape colonies (ECs) during 30 days or more of coculture or cytokine exposure, 10 cm cell culture dishes harboring colonies were washed 3x with PBS and PBS was removed from plates after the last wash step leaving semi-dry dishes. A p10 pipette tip loaded with ∼5 µL of trypsin was used to scratch the center of a colony, detached cells were collected by pipetting, and transferred to a single well of a 12 well cell culture plate. Microscopy images were captured before and after to confirm successful colony isolation. Isolated EC cells were expanded for ∼1 month without CTL or cytokine exposure. The resulting populations were considered regrown ECs.

### Cancer cell viability assays

Cancer cell viability was measured with the CellTiter-Glo Luminescent Cell Viability Assay (Promega #G7571). All coculture cell viability experiments were performed in 12 well culture plates and wells were washed 3x with RPMI media to remove residual CTLs prior to addition of CellTiter-Glo Reagent.

### Crystal violet staining and escape colony quantification

Crystal violet staining was performed in 10 cm cell culture dishes after coculture or cytokine exposure. Cells were washed 3x with 10 mL of PBS, fixed in 5 mL of 90% methanol at −20°C for 10 minutes, and stained in 2 mL of crystal violet solution (Alfa Aesar #B21932, 0.5% crystal violet in 25% methanol) at room temperature for 30 minutes with gentle rocking. Crystal violet solution was removed, dishes were washed 3x with ∼10 mL of distilled water, and placed upside down to dry overnight. Crystal violet staining was imaged using an Interscience Scan 1200 instrument. Cell colonies were counted using the Scan 1200 software (Version 8.0.20 v2.2) with custom settings (Settings = “Dark Colony”, Sensitivity = 72, Discard Debris 1.00 mm) that produced similar results to manual counting.

### Western blot analysis

Protein was extracted from cells using RIPA Lysis and Extraction Buffer (Thermo #89900) with Halt Protease Inhibitor Cocktail (Thermo #87786) and Halt Phosphatase Inhibitor Cocktail (Thermo #78420). To obtain sufficient protein from persister cell samples, six to ten 10 cm dishes were typically pooled per persister cell replicate. Parental cell density and time in media was matched to persister cells for analysis such that 15,000-30,000 parental cells were plated per 10 cm dish and cultured for 3 days before analysis. Dishes were washed at least 3x with 10 mL of cold PBS. Cells were scraped in 2 mL of cold PBS per 10 cm dish, pooled into a 15 mL conical tube, pelleted at 400g at 4°C for 4 minutes, lysed in 100-200 μL of RIPA Buffer, and incubated on ice for 15 minutes with moderate rocking. Lysates were then sonicated for 3 seconds at 50% amplitude using a Branson Digital Sonifier for 3 rounds. Sonicated lysates were centrifuged at ∼14,000g at 4°C for 15 minutes to pellet cell debris and supernatants were transferred to a new tube. Protein concentrations were measured with the Pierce BCA Protein Assay (Thermo #23225).

Protein samples were prepared for denaturing gel electrophoresis in Bolt LDS Sample Buffer (Thermo #B0007) and Bolt Reducing Agent (Thermo #B0004) at 70°C for 10 minutes. Protein samples were then loaded on 4-12% Bolt Bis-Tris Plus Mini Protein Gels (Thermo #NW04122BOX) and electrophoresis was performed using the Mini Gel Tank system (Thermo #A25977) with Bolt MES SDS Running Buffer (Thermo #B0002) at ∼125 V for 45-60 minutes. Chameleon Duo Pre-stained Protein Ladder (LI-COR #928-60000) was used. Following electrophoresis, proteins were transferred to nitrocellulose membranes (Thermo #IB23002) using the iBlot 2 Dry Blotting System (Thermo #IB21001). Membranes were blocked with 5% BSA in TBST (Tris-buffered saline with 0.05% Tween 20, BioPioneer #MB1038) for 1 hour at room temperature with gentle rocking, washed 3x with TBST, and incubated with primary antibodies (diluted in 1% BSA in TBST) at 4°C overnight with gentle rocking. Following primary antibody incubation, membranes were washed 3x with TBST and incubated with IRDye secondary antibodies in 1% BSA in TBST for 1 hour at room temperature with gentle rocking. Following secondary antibody incubation, membranes were washed 3x with TBST, washed 2x with TBS, and imaged for near-infrared fluorescence detection using the Odyssey Classic Imaging System. In some instances, membranes were stripped with NewBlot Nitro Stripping Buffer (LI-COR #928-40030) for reprobing. Quantification of protein band intensity was performed using ImageStudio software. Uncropped western blot images are presented in **Figure S10**.

Primary antibodies for western blotting: β-Tubulin (Invitrogen #MA5-16308, 1:7000 dilution), NY-ESO-1 (Sigma #N2038, 1:2000), Phospho-STAT1 (Y701) (CST #9167, 1:1000), STAT1 (CST #9176, 1:1000), PD-L1 (CST #13684, 1:1000), IDO-1 (CST #86630, 1:1000), Cleaved Caspase 8 (CST #9496, 1:1000), Cleaved Caspase 7 (CST #9491, 1:1000), Cleaved Caspase 3 (CST #9664, 1:1000), Phospho-Histone H2A.X (Ser139) (CST #9718, 1:1000), Histone H2A.X (CST #2595, 1:1000), ICAD/DFFA (abcam #ab108521, 1:10,000), c-IAP2 (CST #3130, 1:1000), c-IAP1 (CST#7065, 1:1000), XIAP (CST #2045, 1:1000), Livin (CST #5471, 1:1000), Survivin (CST #2808, 1:1000), BFL-1 (CST #14093, 1:1000), MCL-1 (CST #94296, 1:1000), BCL-XL (CST #2764, 1:1000), BCL-2 (CST #15071, 1:1000), BCL-w (CST #2724S, 1:1000), NOXA (CST #14766S, 1:1000), BIM (CST #2933T, 1:1000), PUMA (CST #12450, 1:1000), BID (CST #2002, 1:1000), BAD (CST #9239, 1:1000), BAX (CST #5023, 1:1000), BAK (CST #12105, 1:1000), ATF3 (CST #33593, 1:1000), ATF4 (CST #11815, 1:1000), NRF2 (CST #12721, 1:1000), SLC7A11 (CST #12691, 1:1000), H3K14ac (CST #7627, 1:1000), H3K4me3 (CST #9751, 1:1000), H3K9me3 (CST #13969, 1:1000), H3K27me3 (CST #9733T, 1:1000), Histone H3 (CST #3638, 1:1000), MART-1 (CST #34511, 1:1000), MITF (CST #12590, 1:1000), Phospho-ERK1/2 (CST #9101, 1:1000), ERK1/2 (CST #9102, 1:1000). Secondary antibodies for western blotting (1:10,000 dilution): LI-COR 800CW Goat anti-Rabbit IgG (#926-32211), 680RD Goat anti-Mouse (#925-68070), 800CW Goat anti-Mouse (#925-32210), 680RD Goat anti-Rabbit (#926-68071).

### Flow cytometry analysis

CTLs were collected from the coculture supernatant. Live cancer cells were isolated by rinsing plates thoroughly prior to trypsinization, and trace residual CTLs and dead cancer cells which remained despite rinsing were removed from analysis by flow cytometry gating using anti-CD8 antibody to exclude CTLs and cell viability Ghost Dye 510 stain to exclude dead cells. For cell surface staining, cells were washed in PBS, incubated with Ghost Dye 510 viability dye diluted in PBS for 30 minutes at 4°C, washed in PBS + 1% FBS, incubated with conjugated primary antibodies diluted in PBS + 1% FBS for 30 minutes at 4°C, washed in PBS + 1% FBS, resuspended in 2% PFA diluted in PBS, transferred to Round-Bottom Polystyrene Test Tubes (Corning #352235) and analyzed on a BD FACSCanto RUO Analyzer or BD FACSAria II High Speed Sorter. For intracellular staining, these additional steps were added: following washing away Ghost Dye 510, cells were fixed in 4% PFA diluted in PBS for 10 minutes at room temperature, washed in PBS, permeabilized in 0.3% Triton-X-100 diluted in PBS + 1% FBS for 10 minutes at room temperature, and washed in PBS prior to conjugated primary antibody incubation. Unstained cells without primary antibody were included in each experiment. Spectral compensation was performed with compensation beads (Invitrogen #01-2222-41) incubated with conjugated primary antibodies diluted in PBS + 1% FBS for 30 minutes at 4°C. Flow cytometry data were analyzed using FlowJo. Full gating strategies for flow cytometry experiments are presented in **Figures S11-S19**.

Antibodies and dyes used for flow cytometry: NY-ESO-1 AF647 (CST #66920, 1:50), CD25-PE (Invitrogen #12-0259-42, 1:20), CD8-APC (STEMCELL #60022AZ.1, 1:20), CD8 PE (STEMCELL #60022PE.1, 1:20), HLA-A2 APC (Invitrogen #17-9876-42, 1:20), Granzyme B AF700 (BD Pharmingen #560213, 1:100), Cleaved Caspase 3 AF647 (CST #9602, 1:50), Ghost Dye 510 (VWR #10140-8921), and BV510 Mouse IgG1 Isotype Ctrl (Biolegend #400171) was used to compensate for Ghost Dye 510.

### Caspase 3/7 activity measurements

Caspase 3/7 activity was measured by flow cytometry using NucView 530 Caspase-3 Substrate (Biotium #10408/Sigma #SCT105). Cells were washed with PBS, collected by trypsinization, and incubated with the cell viability Ghost Dye Violet 510 for 30 minutes at 4°C to label dead cells for exclusion from flow cytometry analysis. Cells were then washed with PBS + 1% FBS to remove unbound Ghost Dye and then incubated in 500 μL of 5 μM NucView 530 Caspase-3 Substrate (diluted in PBS + 1% FBS) for 30 minutes at room temperature and analyzed by flow cytometry using a BD FACSAria II High Speed Sorter. To assess the viability of caspase-active persister cells, persister cells were sorted by caspase activity, collected in PBS + 10% FBS + antibiotic-antimycotic, and 500 cells from each caspase-activity group were plated into 12 well plates in RPMI 1640 + 10% FBS + antibiotic-antimycotic. Following overnight incubation, cell viability was measured by CellTiter-Glo for a day 1 viability measurement. Cells were then monitored for 6 days and cell viability was again measured by CellTiter-Glo for a day 6 viability measurement. Drug-persister caspase 3/7 sensor data in **Figure S6C** are reanalyzed from Williams et al.^7^

### Mitochondrial membrane potential measurements

Mitochondrial membrane potential was measured by flow cytometry using JC-1 Dye (Invitrogen #T3168). Cells were treated with 50 μM of the mitochondrial oxidative phosphorylation uncoupler carbonyl cyanide m-chlorophenyl hydrazone (CCCP, abcam #ab141229) for 25 minutes as a positive control. A375 parental, CTL-tolerant persister cells, and CCCP-treated cells were collected by trypsinization, counted, resuspended in RPMI 1640, and incubated with 1.5 μM JC-1 dye for 30 minutes at 37°C. Cells were washed in warm PBS, pelleted, and resuspended in Ghost Dye Red 780 (Tonbo 13-0865-T100) for 15 minutes at room temperature. Cells were washed in PBS + 1% FBS, resuspended in 300 μL PBS, and analyzed on a BD FACSCanto RUO flow cytometer. Compensation beads (Invitrogen #01-2222-41) with FITC and PE antibodies were used to compensate green and red fluorescence by JC-1 dye and APC-CY7 was used to compensate for Ghost Dye Red 780.

### Soluble IFNγ measurements

Media was collected at the indicated timepoints during coculture and centrifuged at 400g for 4 minutes to pellet cells and debris. Supernatant was transferred to a new tube and frozen at −80°C. Supernatants were thawed and IFNγ concentrations were measured using Human IFN gamma ELISA Kit (Invitrogen #KHC4021) per product protocol.

### Reverse transcription quantitative real-time PCR (RT-qPCR)

Total RNA was isolated from cells with TRIzol Reagent (Invitrogen #15596026). RNA yield was measured with a NanoDrop spectrophotometer. cDNA was synthesized from 1 μg of RNA by reverse transcription with the RevertAid First Strand cDNA Synthesis Kit (Thermo Scientific #K1622) using an Oligo (dT)_18_ primer. cDNA was diluted 1:1 with nuclease-free water and 2 μL of the diluted cDNA template was used for quantitative real-time PCR using TaqMan Fast Advanced Master Mix (Thermo #4444556) in 20 μL reactions. qPCR reactions were performed using the Bio-Rad CFX96 Real-Time PCR System as follows: 50°C (2 minutes), 95°C (20 seconds), 45 cycles of 95°C (3 seconds) denaturation and 60°C (30 seconds) annealing/extension. Measurements were made using human IFNB1 and GAPDH TaqMan Gene Expression Assays (Thermo #4331182). Relative gene expression for each sample was calculated with the 2^-ΔΔCT^ method using GAPDH as the normalizer gene.

### Single cell RNA-sequencing (scRNA-seq)

A375 parental and CTL-tolerant persister cells were isolated by trypsinization and libraries were generated using the 10X Chromium Single Cell 3 v3 kit (10X Genomics). Quality control of the libraries was conducted with the High Sensitivity D1000 ScreenTape kit with the Agilent TapeStation and then sequenced using a NovaSeq S4 flowcell on an Illumina NovaSeq 6000 sequencer at the UCSD Institute for Genomic Medicine. Data were processed through the Cell Ranger workflow, producing 10X scRNA-seq datasets. A375 drug-tolerant persister and matched parental cell scRNA-seq datasets were previously generated by our lab (Williams A. F. et al., in revision,^7^ accession number: GSE196018).

ScRNA-seq analyses were performed using the Seurat v5 R package.^104^ Each 10X dataset was used to create a unique molecular identified (UMI) count matrix (‘Read10X’) that was then used to create a Seurat object (‘CreateSeuratObject’). Seurat objects were merged and a standard preprocessing workflow without additional dataset integration algorithms was applied. We excluded cells with a low number of detected genes (< 1000), unusually high number of genes (> 7500), or high mitochondrial counts (> 20%). The few detected CD8^+^ T cells (n = 177) within the CTL-tolerant persister cell dataset were removed from analysis by excluding cells with expression of either CD8A, CD3D, CD3E, CD3G, or IL2RA. After filtering, there remained gene expression profiles of 6,348 CTL-tolerant persister cells and 10,526 parental cells from this study, and 4,790 drug-tolerant persister cells and 10,514 parental cells from Williams et al.^7^ Merged and filtered datasets were then sctransform normalized (‘SCTransform’), PCA linear dimensionality reduction was performed (‘RunPCA’), nearest neighbors were computed (‘FindNeighbors’ dims = 1:30), clusters were determined (‘FindClusters’), and UMAP non-linear dimensional reduction was performed (‘RunUMAP’ dims = 1:30). Resulting transcriptional UMAP plots of merged parental and persister cell scRNA-seq datasets were visualized using the Seurat ‘FeaturePlot’ function. To predict the cell cycle status of cells within scRNA-seq datasets, the Seurat ‘CellCycleScoring’ function was applied with obsolete gene names MLF1IP, FAM64A and HN1 updated to their current names CENPU, PIMREG and JPT1, respectively. Cell cycle regression did not noticeably alter cell clustering and we therefore performed data normalization without cell cycle regression.

### Signature scoring

The top 50 genes ranked by adjusted P value in each melanoma cell state signature,^35^ complete ITH MP signatures (∼50 genes each),^61^ and an anastasis signature (41 genes)^40^ were used as gene sets for signature scoring. Genes within signatures analyzed in this study are provided in **Table S1**. Expression of a signature was determined using the Seurat ‘AddModuleScore’ function and Mann-Whitney nonparametric tests were utilized to compare signature scores between groups. Data in **Figure S7B-C** are presented as change in signature score whereby persister cell scores are subtracted by the average score of their experiment-matched parental population. Values above 0 indicate persister cell upregulation of the signature and values below 0 indicate persister cell downregulation of the signature compared to their parental population.

### Gene set enrichment analysis (GSEA) of scRNA-seq data

Preranked gene lists were generated from differentially expressed genes identified using the Seurat function ‘FindMarkers’. Genes that were detected in > 1% of cells by scRNA-seq were included and, to mitigate effects of sparsely detected genes in scRNA-seq data, log2 fold change (log2FC) values were multiplied by the percent detection of each gene in the comparison group the gene is enriched in. Preranked gene lists generated using this population-weighting adjustment approach, as well as ranking solely using log2FC, are shown in **Table S2**. The population-weighted preranked gene lists CTL-persister vs parental (13,218 genes), Drug-persister vs parental (15,233 genes), and CTL-persister vs Drug-persister (15,219 genes) were used for GSEA of scRNA-seq data using GSEA v4.3.2 with 1000 permutati ns.^105,106^ The analyzed gene sets are provided in **Table S1**. Only gene sets with > 70% of genes detected were considered. In **Figure S7D**, significantly enriched gene sets were identified as those with nominal P value < 0.01 and FDR < 0.25.

### Identification of marker genes of persister cell clusters

The Seurat function ‘FindAllMarkers’ was used to identify marker genes of the 20 clusters identified within the parental and persister cell UMAP and we focused on the 7 clusters within persister cell populations (Figures 4D and **S7E**). Differentially expressed genes between clusters were identified by Wilcoxon Rank Sum tests and ROC analysis to generate a ‘predictive power’ ranked list of candidate differentially expressed genes. Only genes detected in > 1% of cells and which show > 0.1 log2FC were included for analysis. Identified cluster marker genes are provided in **Table S3**.

### Whole exome sequencing and variant calling

We performed whole exome sequencing of the clonally-derived A375-A parental reference population and five A375-A-derived ECs that emerged during one month of coculture. Individually isolated ECs were expanded without CTLs prior to sequencing. Approximately 5 million cells from each population were collected by trypsinization, pelleted, snap frozen in liquid nitrogen, stored at −80°C, and sent to Novogene for DNA extraction, library preparation (Agilent SureSelect V6 58M), and Illumina NovaSeq 6000 PE150 sequencing. Each sample was analyzed according to the GATK best practices workflow for data preprocessing for variant discovery followed by the somatic short variant discovery workflow.^107–109^ GATK version 4.2.4.0 pipelines were used. Fastq files were mapped to the hg38 reference genome provided in the GATK shared resource bucket with BWA. Duplicate reads were marked with Picard, and base scores were recalibrated with the GATK tool ‘BaseRecalibrator’. Mutect2 was used to call mutations between the ECs and the reference A375-A parental population. Sequencing errors and contamination were estimated with ‘GetPileUpSummaries’, ‘CalculateContamination’, and ‘FilterMutectCalls’. Variants were annotated with ‘Funcotator’. Low frequency (frequency < 0.3) mutations, which could arise post-treatment during CTL-free expansion of isolated single ECs prior to sequencing, were excluded from the analysis. Output files were analyzed with the Maftools R Bioconductor package version 2.6.05. Mutations with an allele frequency > 0.3 and TLOD6 were called as acquired mutations and are listed in **Table S4**.

### Reexpression of NY-ESO-1

Lentiviral vectors with EF1α-driven human NY-ESO-1 encoding gene CTAG1B (NM_001327.3) and mPGK-driven puromycin resistance gene (Vector ID VB220617-1296ayr), and a control vector with CMV-driven mCherry and EF1α-driven puromycin resistance gene (Vector ID VB900088-2695dmk) were generated by VectorBuilder. E. coli stocks harboring the plasmids were streaked onto LB plates containing 100 μg/mL ampicillin, colonies were isolated and expanded into 100 mL of LB. Plasmid DNA was purified using the Qiagen HiSpeed Plasmid Midi Kit (Qiagen #12643) and measured with a NanoDrop spectrophotometer. To generate lentiviral supernatant, ∼3,800,000 HEK293T cells were plated in 10 cm dishes in DMEM (Gibco #11965118) supplemented with 10% FBS and antibiotic-antimycotic. The next day, HEK293T cells were exposed to a transfection mix containing packaging plasmids psPAX2 (Addgene #12260) and pMD2G (Addgene, #12259), and transfer plasmids in Opti-MEM (Thermo #31985062) and polyethylenimine (PEI, Sigma-Aldrich, #764604) at a 1:3 DNA:PEI ratio. Media was replenished the following day and lentiviral supernatant was collected 48 hours later, centrifuged, filtered through a 0.45 μM filter (Sigma-Aldrich, #SE1M003M00), snap frozen in liquid nitrogen, and stored at −80°C. To transduce A375 cells, 50,000 A375 cells were reverse transduced in 6 well cell culture plates at varying lentiviral supernatant dilutions (0, 1:5, 1:10, 1:50, 1:100, 1:500) for 48 hours. Cells were selected with 2 μg/mL puromycin for 6 days, with puromycin replenished every 2 days. Surviving cells were expanded and NY-ESO-1 expression was measured by western blot.

### Additional chemicals and drugs

Human IFN-gamma antibody (R&D/Fisher #MAB285-SP), human TNF antibody (R&D/Fisher #AF-210-SP), human Fas Ligand antibody (R&D/Fisher #MAB126-SP), concanamycin A (abcam #ab144227), ruxolitinib (MedChemExpress #HY-50856), epacadostat (MedChemExpress #HY-15689), L-tryptophan (Sigma/Neta #T0254), pembrolizumab (anti-PD-1) (Selleckchem #A2005), durvalumab (anti-PDL1) (Selleckchem #A2013), staurosporine (Selleckchem #S1421), quinoline-val-asp-difluorophenoxymethylketone (QVD) (Selleckchem #S7311), LCL161 (MedChemExpress #HY-15518), ferrostatin-1 (APExBIO #A4371), RSL3 (Sigma #SML2234-5MG), panobinostat (Cayman #13280), trichostatin A (Selleckchem #S1045), vorinostat (Selleckchem #S1047), decitabine (5-Aza-2′-Deoxycytidine, Sigma #189826), abemaciclib (CDK4/6 inhibitor) (MedChemExpress #HY-16297).

### Patient tumor slice culture

Melanoma tissue was acquired in accordance with University of California San Diego Institutional Review Board approved protocol #161227. The deidentified sample was a stage IIC primary cutaneous melanoma with BRAF exon 11 p.P453T variant of unknown significance which was surgically resected from an 86 year old male patient and placed into RPMI 1640 media (Gibco #11875119) briefly before being manually cut into 2 mm thick slices. One tumor slice was immediately formalin fixed (10% formalin, phosphate buffered, Newcomer Supply #1090N) for use as a pretreatment control for IHC. The remaining tumor slices were cultured at 37°C with 5% CO_2_ in RPMI 1640 media supplemented with 10% FBS and 1% antibiotic-antimycotic solution (Gibco #15240062). Tumor slices were treated with 10 µg/mL Pembrolizumab (anti-PD-1, Selleck Chemicals #A2005), 10 ng/mL recombinant human IFNγ (Peprotech #300-02), and 10 ng/mL recombinant human TNF (Peprotech #300-01A), and another slice was treated with 500 nM staurosporine (Selleckchem #S1421), for 6 days respectively with media refreshed on day 3.

Following treatment, tumor slices were either formalin fixed and processed for IHC, or were prepared for live cell flow cytometry. For flow cytometry, tumor slices were digested by shaking at 37°C for 25 minutes in RPMI containing collagenase and hyaluronidase (1:10, STEMCELL #07912) and DNAse I (3:20, STEMCELL #07900), followed by manual dissociation with a wide bore 1000 µl pipette tip. The digested tumor tissue was placed through a 70 μM nylon mesh strainer (STEMCELL #27260) and cells were pelleted and incubated with ammonium chloride (STEMCELL #07800) for 5 minutes. The isolated cells were treated with dead cell stain Ghost Dye 510 (Tonbo Biosciences #13-0870) for 30 minutes at 4°C, followed by NucView 530 Caspase-3 Substrate (Biotium #10408/Sigma #SCT105) as described in the flow cytometry methods section. Live caspase-high and caspase-low cell populations were sorted using a BD FACSAria II High Speed Sorter and replated in RPMI 1640 media supplemented with 10% FBS and 1% antibiotic-antimycotic solution for 24 hours. Following this, the fraction of live cells was assessed by combining trypsinated cells together with supernatant containing any live or dead cells, and this pooled cell population was stained with Ghost Dye 510 for 30 minutes at 4°C. The cells were then fixed and permeabilized as described in the flow cytometry methods section before being stained with tumor cell-specific S100B-AF647 antibody (1:50, Abcam #ab196175) for 1 hour at 4°C. Cells were then analyzed by flow cytometry using a BD FACSCanto RUO Analyzer and data were analyzed using FlowJo.

### Mouse tumor experiments

All the animal studies were approved by the University of California San Diego (UCSD) Institutional Animal Care and Use Committee protocol #S15195, and all experiments adhered with all relevant ethical regulations for animal testing and research. Mice were obtained from Charles River Laboratories (Worcester, MA) and housed in individually ventilated micro-isolator cages supplied with acidified water and fed 5053 Irradiated Rodent Diet 20 (Picolab). Temperature for laboratory mice in this facility is maintained between 18–23°C with 40–60% humidity. The vivarium is maintained in a 12 hour light/dark cycle. All animal manipulation activities are conducted in the laminar flow hoods. For orthotropic implantation, 4MOSC1 cells were transplanted (1 million cells per mouse) into the tongue of female C57Bl/6 mice (4–6 weeks of age). Upon tumor formation on day 5, the mice were randomized into groups. For drug treatment, the mice were treated by intraperitoneal injection with 10 mg/kg anti-PD-1 antibody (BioXCell, Clone J43, #BE0033-2) on day 6 and day 8 post-implantation. Mice were sacrificed on day 10 and tumors were harvested and formalin fixed for IHC analysis. Tumors which decreased in volume by at least 40% from their maximum volume during treatment were considered responders and tumors which did not decrease in volume during treatment were considered nonresponders.

### Immunohistochemistry staining and analysis

Formalin-fixed tumor slices were paraffin-embedded and sectioned for IHC staining by Histoserv (Germantown, Maryland, USA) and the UCSD Tissue Technology Shared Resource. Primary antibodies against ATF3 (1:100, Cell Signaling Technology #18665), mouse IDO1 (1:200, LS-Bio #LS-B13059), human IDO1 (1:200, Cell Signaling Technology #86630), S100B (1:500, Abcam #ab52642), and pan-Cytokeratin (1:1000, Dako #Z0622) were used. Antigen retrieval was performed using a citrate-based antigen unmasking solution (Vector Laboratories #H-3300) at 95°C for 30 minutes. Staining was performed using an Intellipath automated IHC Stainer (BioCare Medical, Irvine, USA) and primary antibodies were incubated for 1 hour at room temperature. Primary antibodies against ATF3, IDO1, S100B, and pan-Cytokeratin were labeled with Anti-Rabbit UltraPolymer HRP (Cell IDx #2RH). Secondary antibodies were visualized using 3,3’-Diaminobenzidine Solutions Kit (VWR #95041-478) and counterstained with Mayer’s Hematoxylin (Sigma #51275).

The slides were scanned using an Aperio AT2 – Digital whole Slide Scanner (Aperio Group, Sausalito, USA) and images were captured using Aperio ImageScope software (version 12.4.6). To quantify markers, five equivalently sized fields were randomly collected per tumor by blinded researchers. To quantify the relative area of cells with positive staining for each IHC marker, analysis was performed using Fiji software^110^ with a color deconvolution plug-in installed^111,112^ which unmixed the signal from the stains and allowed for area quantification of the 3,3’-diaminobenzidine (DAB) chromogen signal within each image. A pixel threshold was chosen for each IHC marker to differentiate apparent signal from background signal and this threshold was equally applied to all images within each blinded group to calculate the relative DAB positive percentage area of cells.

### Statistical analyses

No statistical methods were used to predetermine sample size. Unless otherwise stated, statistical tests and graphing were performed with GraphPad Prism v.10.1.1 and P values were calculated using unpaired, two-tailed unpaired t-tests assuming unequal variance.

### Data and code availability

ScRNA-seq and whole exome sequencing data are available upon request and will be accessible in the NCBI Genome Expression Omnibus (GEO) and NCBI Sequence Read Archive upon final publication of this manuscript.

## Acknowledgements

We thank the Flow Cytometry Core at the San Diego Center for AIDS Research (SD CFAR), an NIH-funded program (P30 AI036214), which is supported by the following NIH Institutes and Centers: NIAID, NCI, NHLBI, NIA, NICHD, NIDA, NIDCR, NIDDK, NIMH, NIMHD, NINR, FIC, and OAR. We further thank the UCSD Tissue Technology Shared Resource for technical assistance, Alex Wenzel for advice on scRNA-seq GSEA, and Susan Kaech and Jack Bui for useful discussions. BioRender.com was used to generate figure cartoons. This research was supported by NIH training grants T32CA067754 (M.X.W. and A.F.W.), T32GM007752 (B.E.M), and T32CA009523 (A.F.W.), and DOD CDMRP MRP Idea Award HT9425-23-1-0719, DOD CDMRP MRP Melanoma Academy Scholar Award HT9425-24-1-0288, a Curebound Discovery Grant, sponsored research grant BG104446 from BridgeBio subsidiary Ferro Therapeutics, American Cancer Society IRG #19-230-48, UCSD Moores Cancer Center Specialized Cancer Center Support Grant NIH/NCI P30CA023100, NIH/NCI R01CA212767, V Foundation for Cancer Research V Scholar Award V2021-035, Bristol-Myers Squibb Melanoma Research Alliance Young Investigator Award, University of California Research Initiatives Cancer Research Coordinating Committee Seed Grant C23CR5537, University of California Academic Senate Bridge Grant BG104446, Tower Cancer Research Foundation Career Development Grant, and The Skin Cancer Foundation Ashley Trenner Research Grant Award (M.J.H.).

## Author contributions

M.X.W., B.E.M., and M.J.H. conceived and supervised the study and wrote the manuscript, M.X.W., B.E.M., T.B-P., F.A.H., S.H.H., C.P.L., C.E.T., M.H.P., D.A.G.G., T.T., A.E.S. performed the experiments and analyzed the data, B.S.Y., B.E.M., and J.S.G. performed and supervised the mouse studies, M.X.W., and A.F.W. performed bioinformatic analyses, T.G., G.A.D., and S.J.P. provided patient melanoma tissue.

## Declaration of interests

M.J.H. is a cofounder, consultant, and research funding recipient of BridgeBio subsidiary Ferro Therapeutics.

## Supplemental information

Document S1. Figures S1-S9

Document S2. Figure S10 (uncropped western blot images)

Document S3. Figures S11-S19 (gating strategies for flow cytometry experiments)

Table S1. Excel file containing gene sets analyzed by scRNA-seq

Table S2. Excel file containing differentially expressed gene lists used for GSEA

Table S3. Excel file containing marker genes of cell clusters identified by scRNA-seq

Table S4. Excel file containing mutations identified within ECs isolated from A375-A

