## Supplementary material for "Antigenic cancer persister cells survive direct T cell attack": Document S1. Figures S1-S9

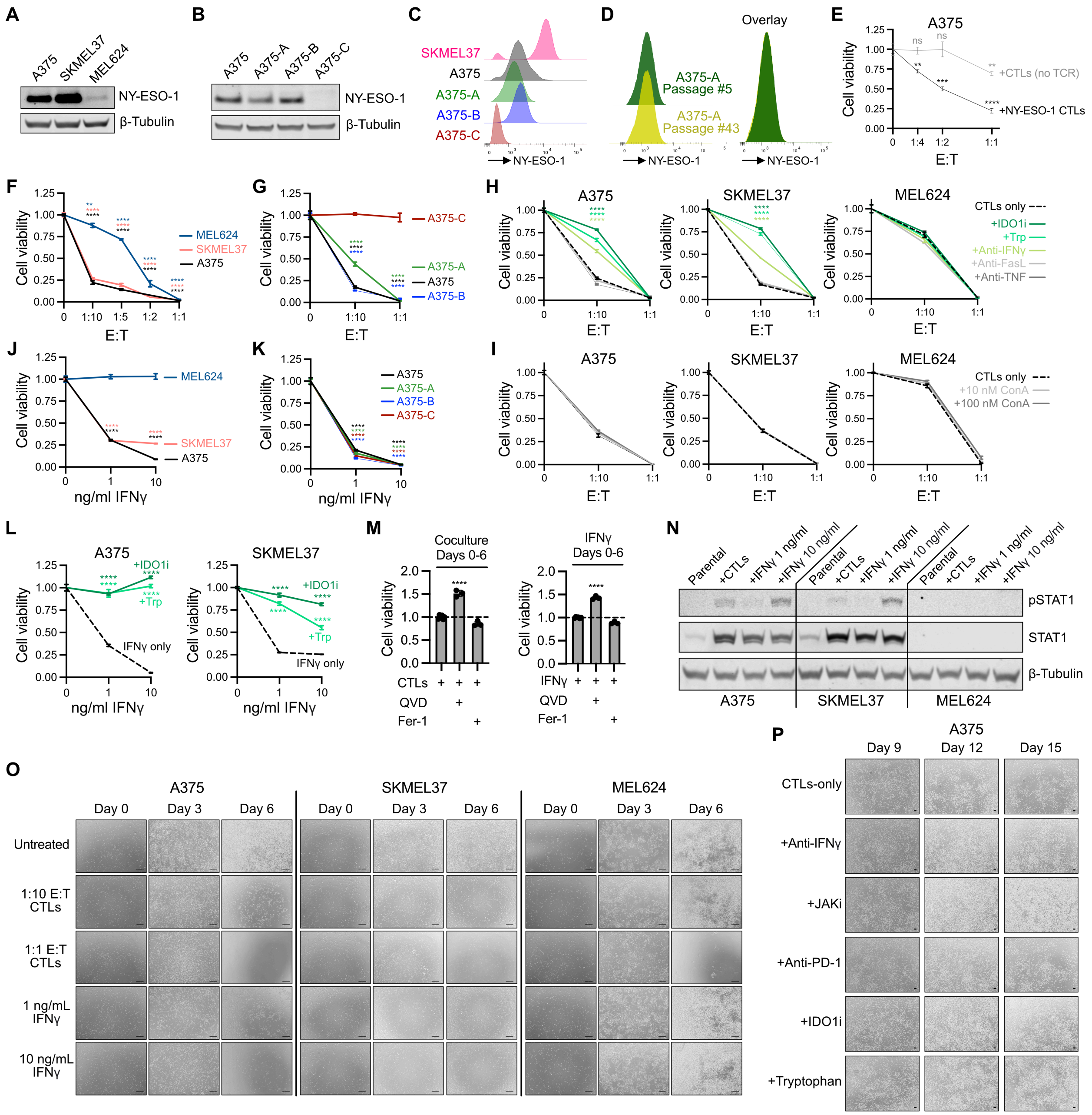

**Figure S1. Characterization of NY-ESO-1 CTL coculture models. (A-B)** Western blot and **(C)** flow cytometry analysis of NY-ESO-1 levels. **(D)** A375 subclone NY-ESO-1 expression levels are stable during prolonged cell culture without CTLs. **(E)** CTLs require transduced NY-ESO-1 TCR to kill target cells at CTL$^{low}$ E:Ts < 1:1 (3 days, t-tests versus untreated cells). **(F-L)** Cell viability after 6 days of **(F-I)** coculture with NY-ESO-1 CTLs or **(J-L)** recombinant IFNγ exposure. **(F-G)** CTL and tumor cell coculture (t-tests versus untreated cells). **(H-I)** Coculture with added **(H)** neutralizing IFNγ antibody (1 μg/mL), TNF antibody (1 μg/mL), FasL antibody (1 μg/mL), and IDO1 inhibitor (2 μM epacadostat) or supplementation with tryptophan (100 μg/mL) (t-tests versus CTLs only coculture) and **(I)** perforin inhibitor concanamycin A (ConA). CTLs were pretreated with 10 nM or 100 nM ConA for 2 hours and washed thoroughly to remove trace ConA prior to coculture. **(J-K)** Recombinant IFNγ exposure alone (t-tests versus untreated cells) and **(L)** with added IDO1 inhibitor (2 μM epacadostat) or tryptophan (100 μg/mL) (t-tests versus IFNγ-only exposures). **(M)** A375 cell viability after 6 days of $CTL^{low}$ coculture or recombinant IFNγ (1 ng/ml) with cotreatment with death pathway inhibitors QVD (10 μM) or ferrostatin-1 (2 μM) (n = 3-6, t-tests versus no inhibitor treatments). **(N)** Western blot of cell lines exposed to $CTL^{low}$ or recombinant IFNγ for 3 days. **(O-P)** Representative microscopy of **(O)** cell line responses during 6 days of NY-ESO-1 CTL coculture or recombinant IFNγ exposure (scalebars, 500 μm) and **(P)** A375 cells co-treated with the indicated treatments from days 9-15 of NY-ESO-1 CTL coculture (scalebars, 100 μm). N = 3, mean ± SD are plotted, and two-tailed unpaired t-tests were performed unless stated otherwise. ns $P > 0.05$; *$P < 0.05$; **$P < 0.01$; ***$P < 0.001$; ****$P < 0.0001$.

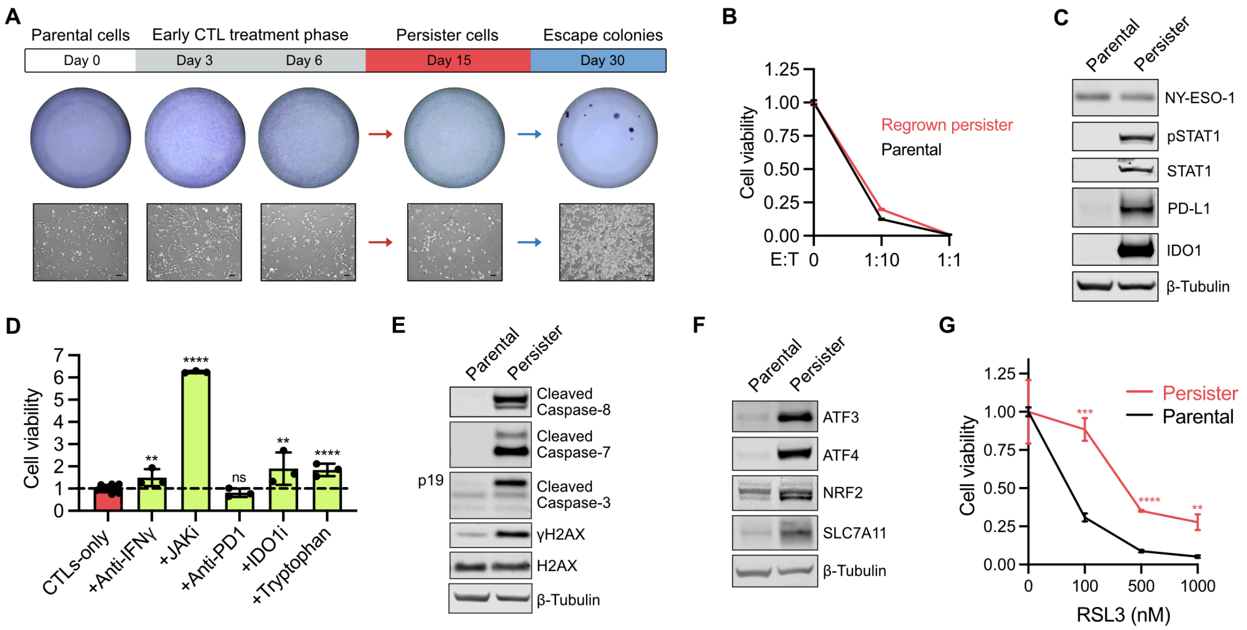


**Figure S2. Characterization of SKMEL37 CTL-tolerant persister cells. (A)** Representative crystal violet staining and microscopy depicting SKMEL37 melanoma cells during 1 month of NY-ESO-1 CTL$^{low}$ coculture. Scale bars, 100 µm. **(B)** Cell viability of SKMEL37 parental cells and CTL-tolerant persister cells which had been allowed to regrow for one month without CTLs prior to challenge with 6 days of NY-ESO-1 CTLs. **(C)** Western blot of NY-ESO-1 and selected IFNγ-signaling nodes within SKMEL37 parental and CTL-tolerant persister cells derived from CTL$^{low}$ coculture. **(D)** SKMEL37 cell viability after 15 days of CTL$^{low}$ coculture with indicated treatments added on days 9-15 of coculture (CTLs-only n = 9, all other conditions n = 3, t-tests versus CTLs-only) with neutralizing IFNγ antibody (1 µg/mL), JAK inhibitor (1 µM ruxolitinib), anti-PD-1 antibody (10 µg/mL pembrolizumab), IDO1 inhibitor (2 µM epacadostat), or tryptophan (100 µg/mL). **(E)** Western blot of caspase cleavage and the DNA damage marker γH2AX in SKMEL37 parental cells and CTL-tolerant persister cells. The caspase 3 partial cleavage p19 fragment is labeled. **(F)** Western blot of stress response factors in SKMEL37 parental cells and CTL-tolerant persister cells. **(G)** SKMEL37 parental cell and CTL-tolerant persister cell viability after 1 day of treatment with GPX4 inhibitor RSL3 (t-test versus parental). N = 3, mean ± SD are plotted, and two-tailed unpaired t-tests were performed unless stated otherwise. ns $P > 0.05$; *$P < 0.05$; **$P < 0.01$; ***$P < 0.001$; ****$P < 0.0001$.

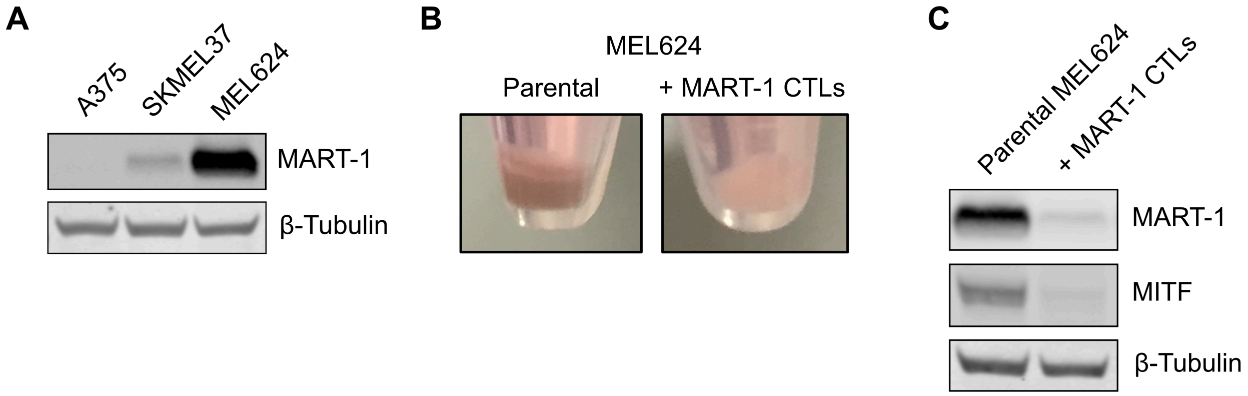


**Figure S3. MEL624 cells dedifferentiate and lose melanocytic MART-1 antigen expression early during MART-1 CTL exposure. (A)** Western blot of endogenous expression of the melanocytic antigen MART-1 in human melanoma cell lines. **(B)** Cell color of pelleted parental MEL624 cells and MEL624 cells after 6 days of MART-1 CTL coculture demonstrating loss of pigmentation from dedifferentiation during coculture. **(C)** Western blot of MART-1 and MITF expression in parental MEL624 cells and MEL624 cells following 6 days of MART-1 CTL coculture.

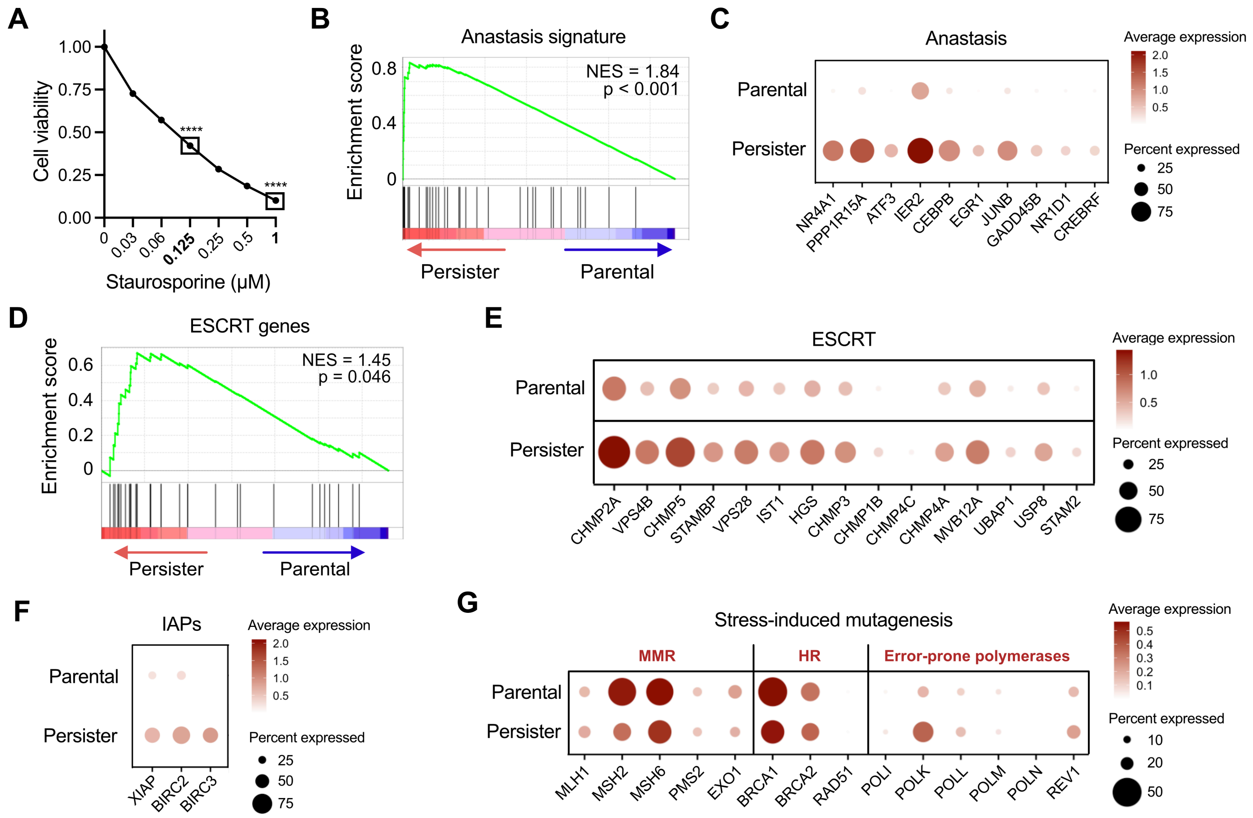


**Figure S4. Additional CTL-tolerant persister cell transcriptional features. (A)** A375 parental cell viability measured 2 days after 4 hour treatment with the apoptosis inducer staurosporine (STS) (n = 3, two-tailed unpaired t-tests versus untreated cells, $****P < 0.0001$). The boxed conditions were used as controls representing moderately lethal (0.125 µM STS) and highly lethal (1 µM STS) apoptotic stress for Figures 2D and 2G. **(B-E)** A375 scRNA-seq GSEA enrichment score and leading edge subset genes for **(B-C)** the anastasis gene set[40] and **(D-E)** ESCRT machinery genes. **(F)** A375 scRNA-seq expression of IAP genes and **(G)** genes previously implicated in stress-induced mutagenesis.[5]

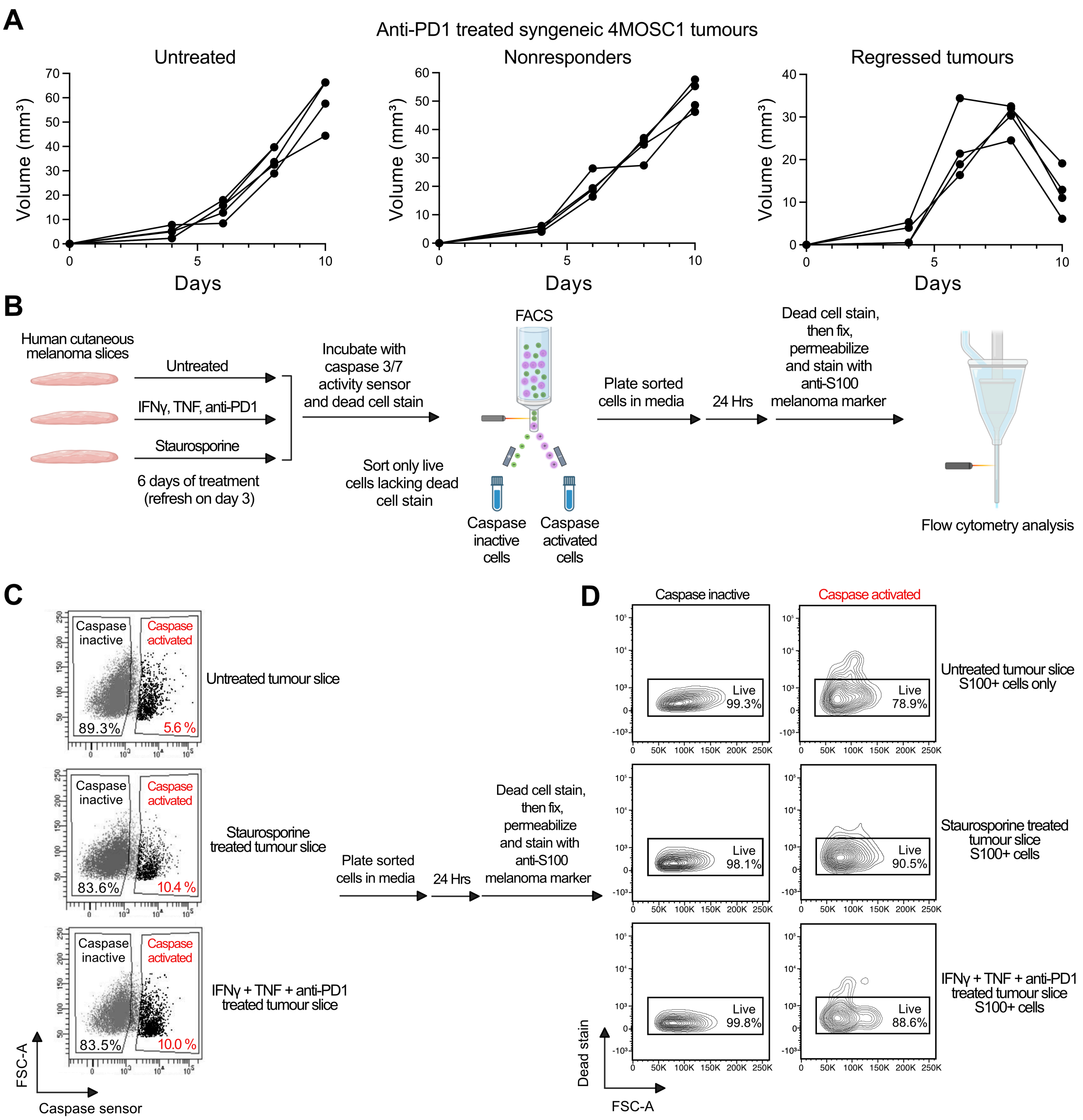


**Figure S5. Analysis of mouse and human tumors treated with immunotherapy. (A)** 4MOSC1 syngeneic tumor-bearing mice categorized based on tumor response during treatment with 10 mg/kg anti-PD-1 (n = 4-5 mice). **(B)** Schematic of human cutaneous melanoma tumor slice treatment and analysis for viable cells with activated caspase 3/7 activity. **(C)** Additional data and representative flow cytometry plots for Figure 3K. Surgically resected primary human cutaneous melanoma tissue treated with 10 µg/mL anti-PD-1, 10 ng/mL IFNγ, and 10 ng/mL TNF, or with apoptosis inducer 500 nM staurosporine, for 6 days in culture were sorted based on caspase activity. Treatment enriched for caspase activated cells which were subsequently sorted, replated and cultured for 24 hours before analysis by flow cytometry to test for viability. **(D)** Analysis of cell viability after sorting for caspase activity and replating for 24 hours. Live cells are indicated by exclusion of dead cell stain Ghost Dye 510. Only S100 positive melanoma cells are shown.

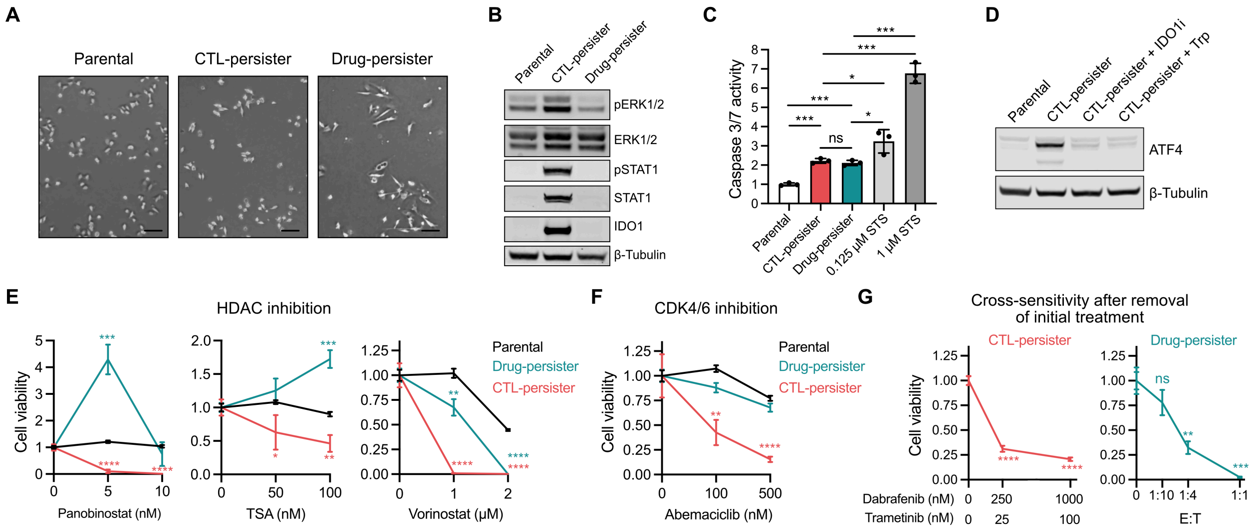


**Figure S6. A375 CTL- and drug-tolerant persister cell comparisons. (A)** Representative microscopy of A375 parental, CTL-persisters and drug-persisters that survive 15 days of NY-ESO-1 CTL$^{low}$ coculture or 250 nM BRAF inhibitor dabrafenib and 25 nM MEK inhibitor trametinib treatment (scalebars, 100 µm). **(B)** Western blot analysis of pERK, pSTAT1, and IDO1 levels in CTL- and drug-persisters. **(C)** Flow cytometry of caspase 3/7 activity. Data are the same as in Figure 2D with added drug-persister caspase activity. **(D)** ATF4 expression in A375 CTL-persisters ± 2 µM IDO1 inhibitor epacadostat or 100 µg/mL tryptophan supplementation (Trp) added during coculture days 12-15. **(E)** Persister cell viability after 15 day cotreatments alongside CTLs or BRAFi and MEKi with HDAC inhibitors panobinostat, trichostatin A (TSA), or vorinostat (n = 3-6, t-tests versus parental). **(F)** Persister cell viability after 3 days of CDK4/6 inhibitor (abemaciclib) treatment (n = 3-6, t-tests versus parental). **(G)** CTL-persister viability after 3 day treatment with dabrafenib and trametinib (left) and drug-persister viability after 3 day coculture with CTLs (right). E:T is based on initially plated cell count prior to any treatment. In contrast to Figure 4J, initial CTL or drug treatments used to derive persister cells were removed and the adherent persister cells were rinsed immediately prior to addition of the alternate treatment (t-tests versus untreated cells). N = 3, mean ± SD are plotted, and two-tailed unpaired t-tests were performed unless stated otherwise. ns $P > 0.05$; *$P < 0.05$; **$P < 0.01$; ***$P < 0.001$; ****$P < 0.0001$.

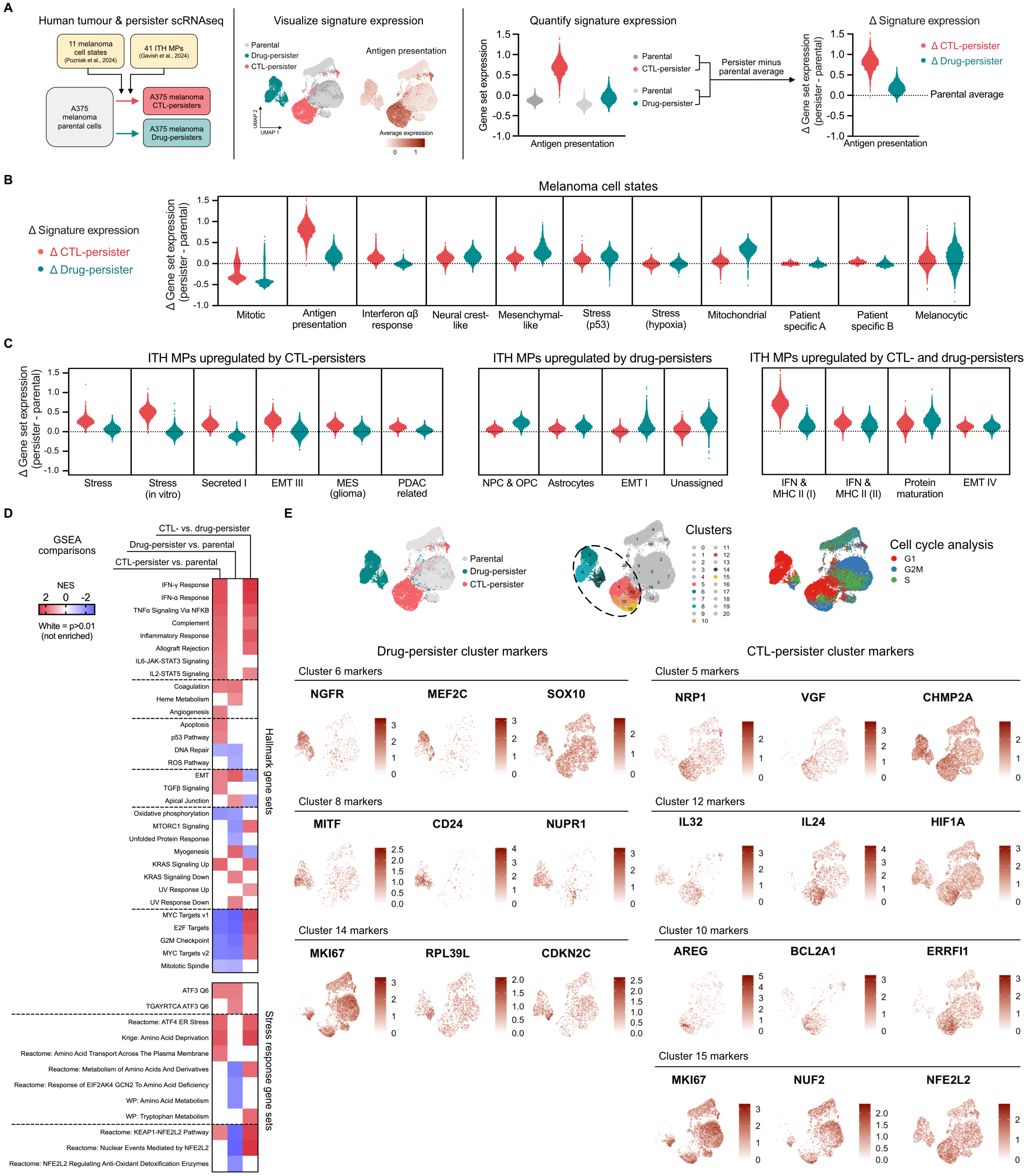

A
Human tumour & persister scRNAseq
11 melanoma cell states (Pozniak et al., 2024)
41 ITH MPs (Gavish et al., 2024)
A375 melanoma parental cells
A375 melanoma CTL-persisters
A375 melanoma Drug-persisters
Visualize signature expression
Parental
Drug-persister
CTL-persister
UMAP 1
UMAP 2
Antigen presentation
Average expression
0
1
Quantify signature expression
Gene set expression
Antigen presentation
Parental
CTL-persister
Parental
Drug-persister
Persister minus parental average
Δ Signature expression
Δ CTL-persister
Δ Drug-persister
Δ Gene set expression (persister - parental)
Parental average
Antigen presentation
B
Melanoma cell states
Δ Signature expression
Δ CTL-persister
Δ Drug-persister
Δ Gene set expression (persister - parental)
Mitotic
Antigen presentation
Interferon αβ response
Neural crest-like
Mesenchymal-like
Stress (p53)
Stress (hypoxia)
Mitochondrial
Patient specific A
Patient specific B
Melanocytic
C
ITH MPs upregulated by CTL-persisters
ITH MPs upregulated by drug-persisters
ITH MPs upregulated by CTL- and drug-persisters
Δ Gene set expression (persister - parental)
Stress
Stress (in vitro)
Secreted I
EMT III
MES (glioma)
PDAC related
NPC & OPC
Astrocytes
EMT I
Unassigned
IFN & MHC II (I)
IFN & MHC II (II)
Protein maturation
EMT IV
D
GSEA comparisons
CTL- vs. drug-persister
Drug-persister vs. parental
CTL-persister vs. parental
NES
2 0 -2
White = p>0.01 (not enriched)
IFN-γ Response
IFN-α Response
TNFα Signaling Via NFKB
Complement
Inflammatory Response
Allograft Rejection
IL6-JAK-STAT3 Signaling
IL2-STAT5 Signaling
Coagulation
Heme Metabolism
Angiogenesis
Apoptosis
p53 Pathway
DNA Repair
ROS Pathway
EMT
TGFβ Signaling
Apical Junction
Oxidative phosphorylation
MTORC1 Signaling
Unfolded Protein Response
Myogenesis
KRAS Signaling Up
KRAS Signaling Down
UV Response Up
UV Response Down
MYC Targets v1
E2F Targets
G2M Checkpoint
MYC Targets v2
Mitotic Spindle
Hallmark gene sets
ATF3 Q6
TGAYRTCA ATF3 Q6
Reactome: ATF4 ER Stress
Krige: Amino Acid Deprivation
Reactome: Amino Acid Transport Across The Plasma Membrane
Reactome: Metabolism of Amino Acids And Derivatives
Reactome: Response of EIF2AK4 GCN2 To Amino Acid Deficiency
WP: Amino Acid Metabolism
WP: Tryptophan Metabolism
Reactome: KEAP1-NFE2L2 Pathway
Reactome: Nuclear Events Mediated by NFE2L2
Reactome: NFE2L2 Regulating Anti-Oxidant Detoxification Enzymes
Stress response gene sets
E
Parental
Drug-persister
CTL-persister
Clusters
Cell cycle analysis
G1
G2M
S
Drug-persister cluster markers
CTL-persister cluster markers
Cluster 6 markers
NGFR
MEF2C
SOX10
Cluster 5 markers
NRP1
VGF
CHMP2A
Cluster 8 markers
MITF
CD24
NUPR1
Cluster 12 markers
IL32
IL24
HIF1A
Cluster 14 markers
MKI67
RPL39L
CDKN2C
Cluster 10 markers
AREG
BCL2A1
ERRFI1
Cluster 15 markers
MKI67
NUF2
NFE2L2

**Figure S7. A375 CTL- and drug-tolerant persister cell transcriptional comparisons. (A)** Pipeline examining gene signatures representing melanoma cell states and intratumor heterogeneity meta-programs (ITH MPs) identified within human tumors (see **Table S1** for gene sets). Panels **B** and **C** are presented as changes in (Δ) signature expression between persisters and their experiment-matched parental population. **(B-C)** All presented comparisons were significant ($P$ < 0.0001, Mann-Whitney tests) except the drug-persister versus parental comparison of the EMT-III ITH MP ($P$ > 0.05) in **C**. **(B)** Analysis of persister expression of melanoma cell state signatures.[35] **(C)** Analysis of persister expression of selected upregulated ITH MPs.[61] All ITH MPs with > 0.10 median persister expression difference versus parental and significant upregulation versus parental by gene set enrichment analysis (GSEA, $P$ < 0.01, not shown) are shown. **(D)** GSEA of Hallmark gene sets and ATF3, ATF4, NRF2, and amino acid stress-related gene sets within the Molecular Signatures Database (MSigDB) using nominal $P$ value < 0.01 and FDR < 0.25 as the significance threshold. Nonsignificant gene sets are indicated in white. **(E)** UMAPs depicting cell cycle stage and expression of selected marker genes that define the persister cell subpopulations which are presented in Figure 4D. Marker genes that define each subpopulation are listed in **Table S3**.

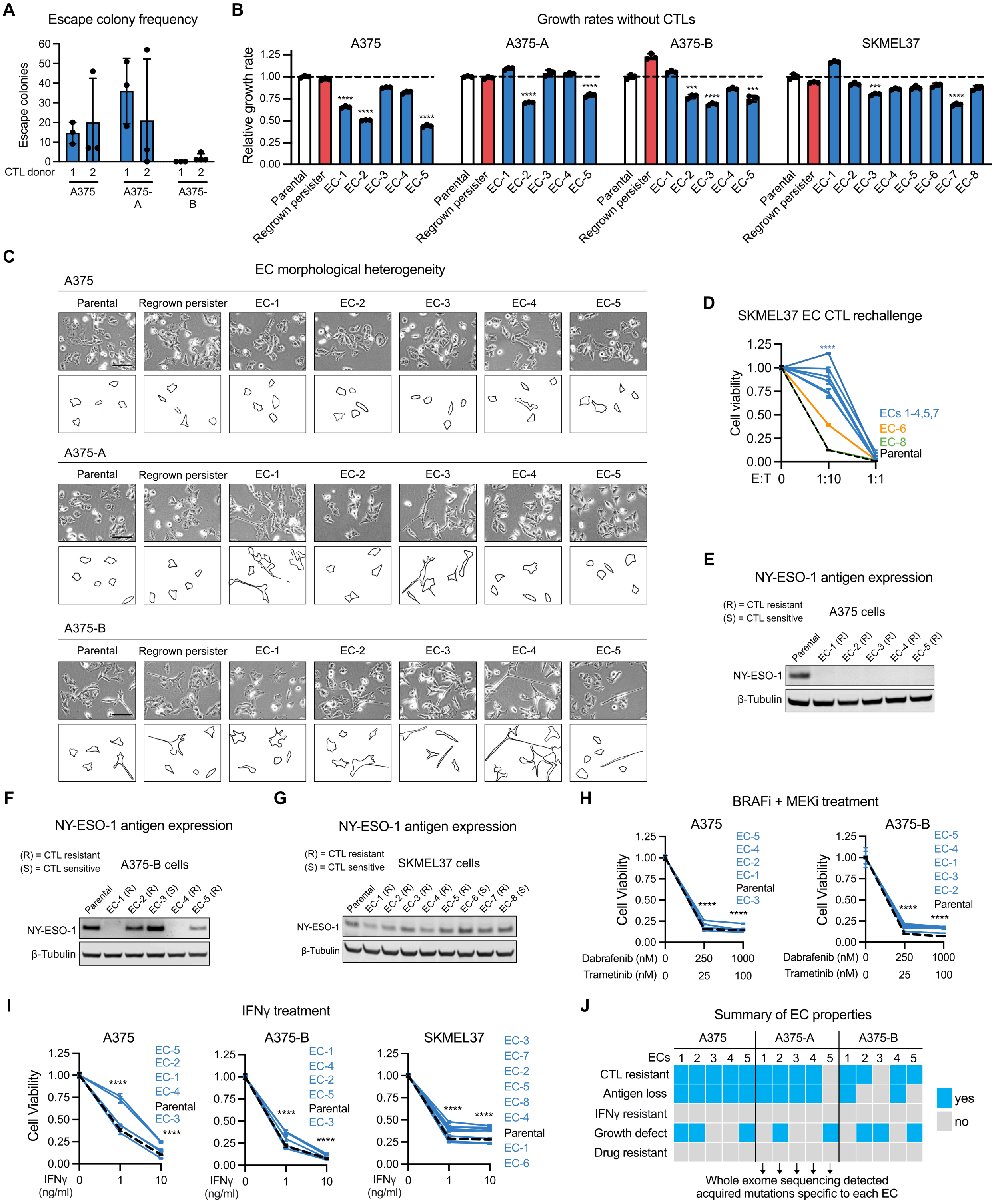


**Figure S8. Characterization of escape colonies isolated during CTL coculture. (A)** Representative quantification of escape colony (EC) formation after 1 month of CTL$^{low}$ coculture using CTLs derived from two different PBMC donors (n = 3-4). **(B)** 3 day CTL-free growth rates of parental, regrown persister, and EC cells each previously allowed to regrow without CTLs for one month (t-tests versus parental). Cells with > 20% decrease in growth rate compared to parental are classified as growth defective in **J**. **(C)** Microscopy of A375 parental, regrown persister, and ECs regrown without CTLs. Traced outlines of selected cell shapes are shown to highlight heterogenous cell morphologies. Scale bars, 100 µm. **(D)** Cell viability after 6 days of NY-ESO-1 CTL coculture of SKMEL37 parental and ECs which had been allowed to regrow without CTLs prior to CTL rechallenge (t-tests for ECs 1-4, 5 and 7 versus parental at 1:10 E:T). **(E-G)** Western blots of NY-ESO-1 expression in parental and escape colony cells derived from $CTL^{low}$ coculture. A375-A data are in Figure 5F. (R) and (S) indicate whether an EC is resistant or sensitive to CTLs after CTL-free regrowth, as measured in **D** and Figure 5B. **(H)** Parental and regrown EC cell viability after 3 days of BRAFi dabrafenib and MEKi trametinib (*****P* for all cell lines and drug concentrations versus untreated cells). A375-A data are in Figure 5K. **(I)** Parental and regrown EC viability after 6 days of recombinant IFNγ exposure. A375-A data are in Figure 5L (n = 3-6, *****P* for all cell lines and IFNγ concentrations versus untreated cells). **(J)** Summary of A375, A375-A, and A375-B EC properties. N = 3, mean ± SD are plotted, and two-tailed unpaired t-tests were performed unless stated otherwise. $*P < 0.05$; $**P < 0.01$; $***P < 0.001$; $****P \leq 0.0004$.

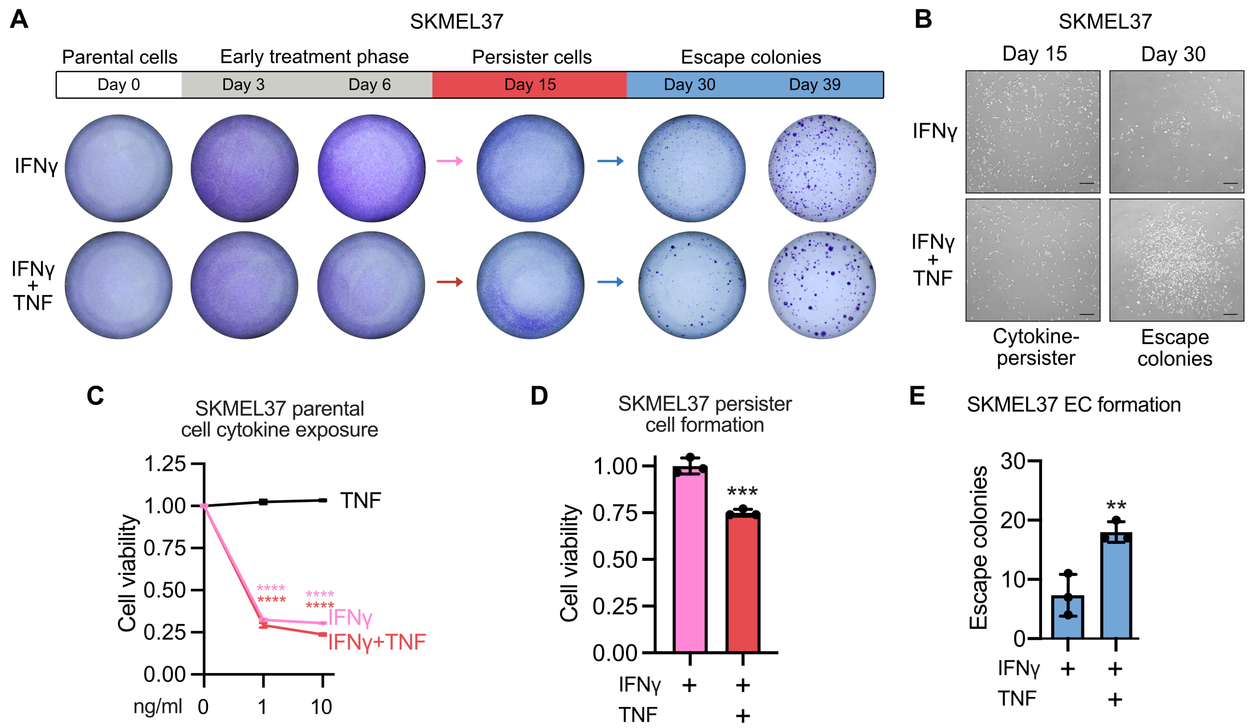


**Figure S9. SKMEL37 cytokine-tolerant persister cells and cytokine escape colonies. (A)** SKMEL37 cells treated with recombinant IFNγ ± TNF exposure (10 ng/ml each). **(B)** Microscopy of initial SKMEL37 escape colony (EC) formation between days 15 and 30 demonstrating TNF promotes earlier ECs. Scalebars, 100 µm. **(C)** SKMEL37 parental cell viability after 6 days of recombinant IFNγ ± TNF exposure (t-tests versus untreated cells). **(D)** SKMEL37 persister cell viability after 15 days of recombinant IFNγ ± TNF exposure. **(E)** Quantification of SKMEL37 EC formation after 39 days of IFNγ ± TNF exposure. N = 3, mean ± SD are plotted, and two-tailed unpaired t-tests were performed unless stated otherwise. *$P < 0.05$; **$P < 0.01$; ***$P < 0.001$; ****$P \leq 0.0001$.
