## Supplementary figures and images for "Antigenic cancer persister cells survive direct T cell attack"

### Document S2. Figure S10

**Figure S10:** Uncropped western blot images.

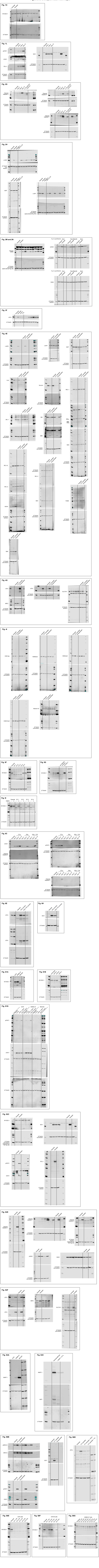
