## Supplementary material for "Antigenic cancer persister cells survive direct T cell attack": Document S3. Figures S11-S19

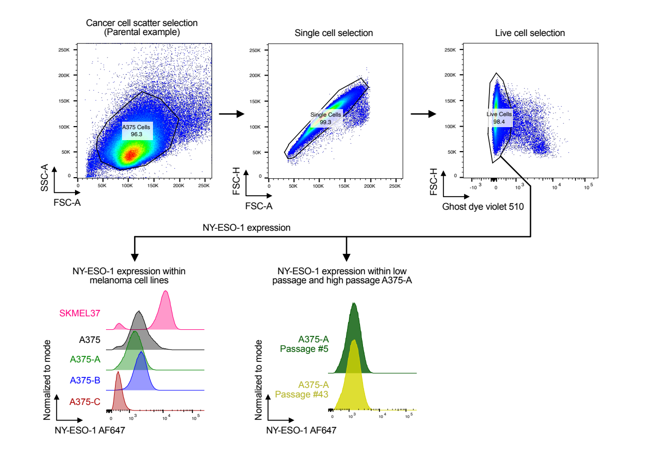


**Figure S11:** Flow cytometry gating strategy for **Figure S1C-D**.

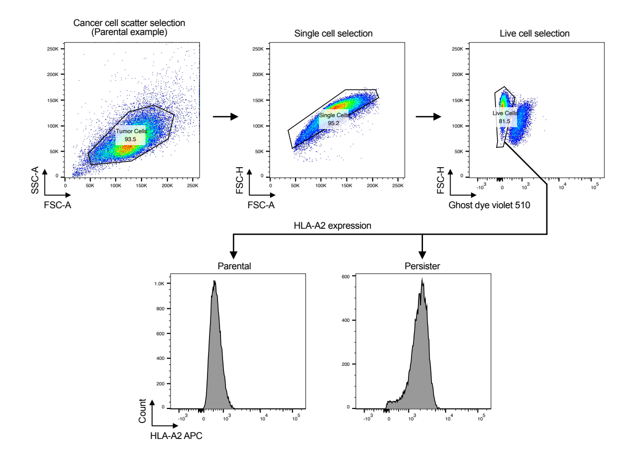


**Figure S12:** Flow cytometry gating strategy for **Figure 1I**.

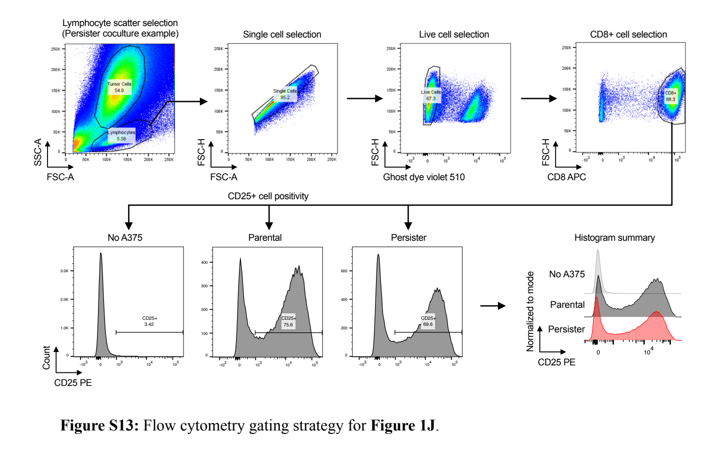


**Figure S13:** Flow cytometry gating strategy for **Figure 1J**.

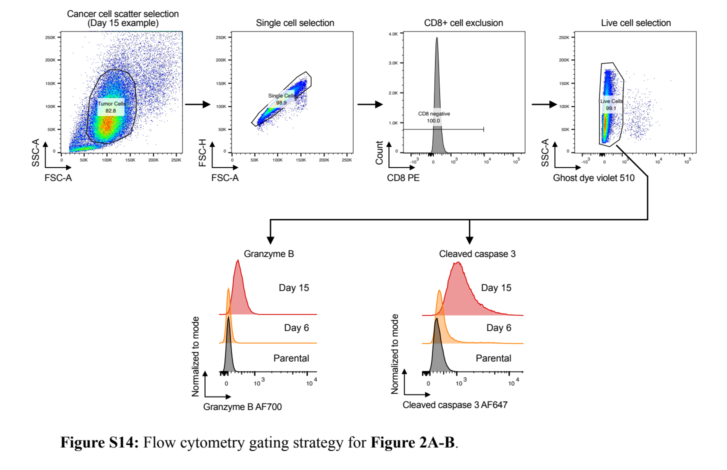


**Figure S14:** Flow cytometry gating strategy for **Figure 2A-B**.

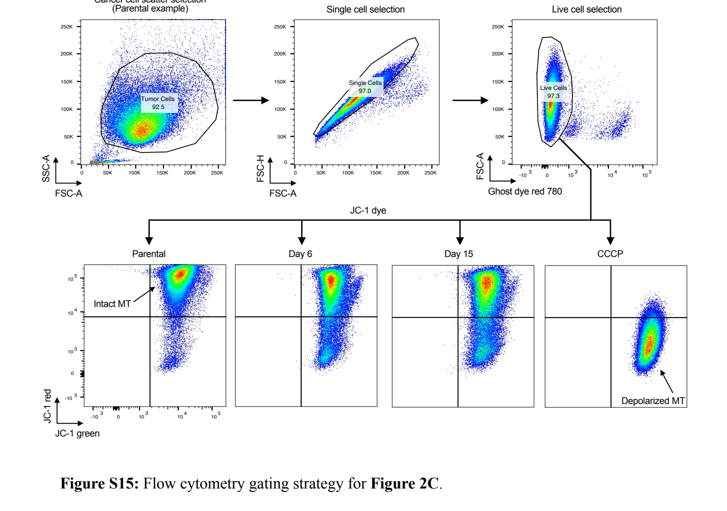


**Figure S15:** Flow cytometry gating strategy for **Figure 2C**.

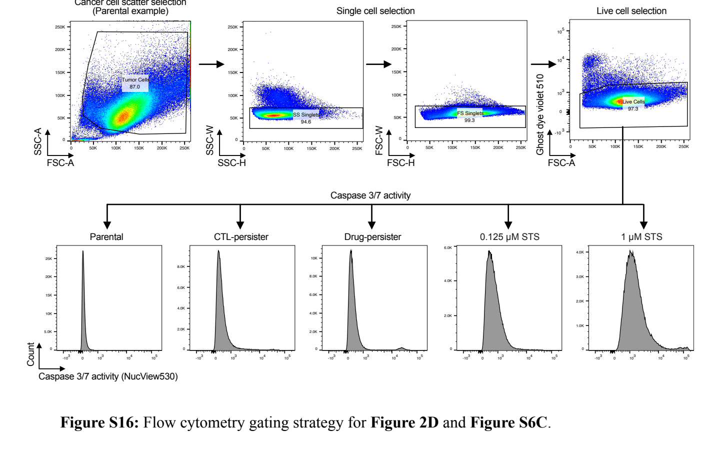


**Figure S16:** Flow cytometry gating strategy for **Figure 2D** and **Figure S6C**.

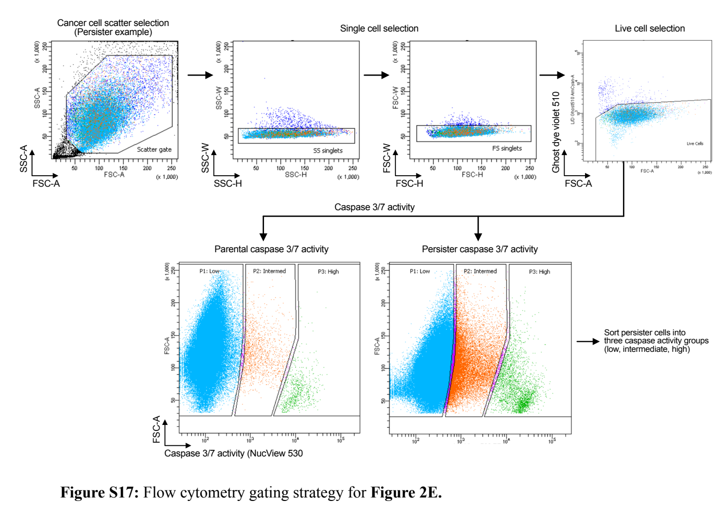


**Figure S17:** Flow cytometry gating strategy for **Figure 2E.**

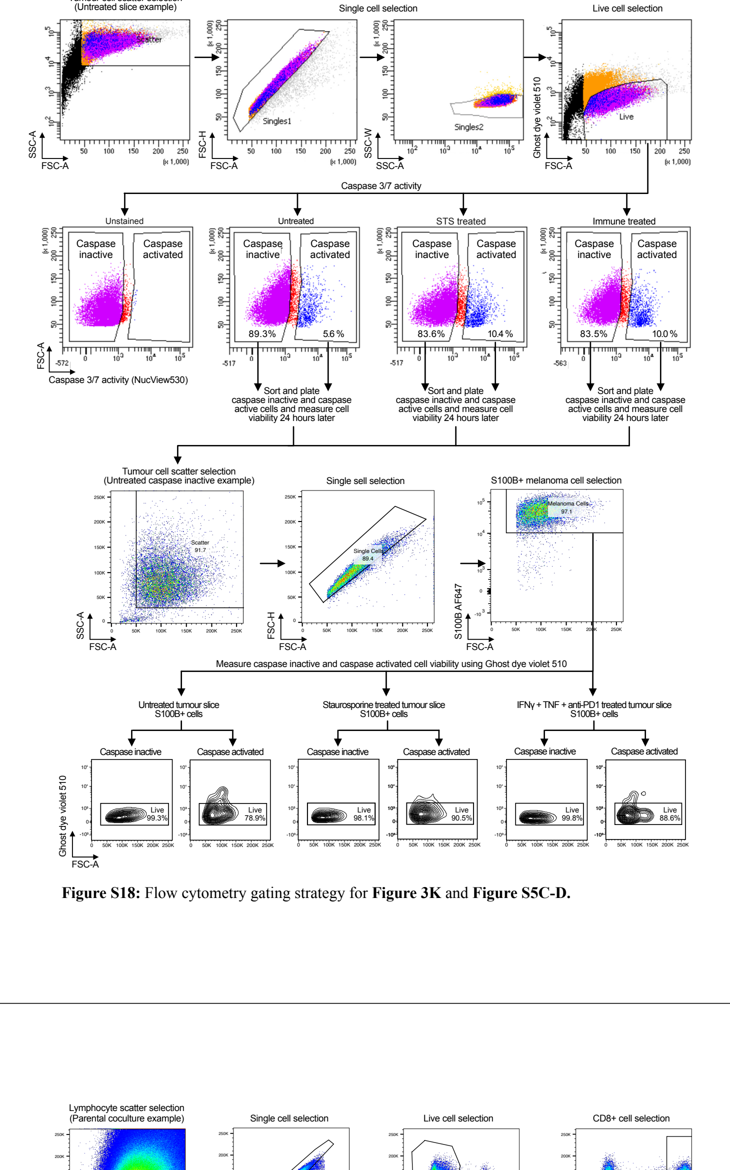


**Figure S18:** Flow cytometry gating strategy for **Figure 3K** and **Figure S5C-D.**

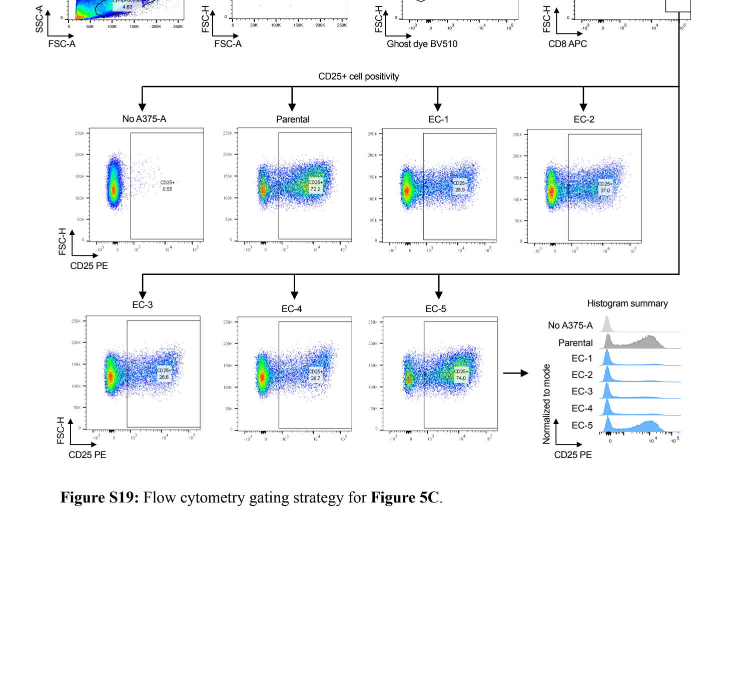


**Figure S19:** Flow cytometry gating strategy for **Figure 5C**.
